# The membrane-cytoplasmic linker defines activity of FtsH proteases in *Pseudomonas aeruginosa* clone C

**DOI:** 10.1101/2023.06.19.545564

**Authors:** Gina D Mawla, Shady Mansour Kamal, Lianying Cao, Pasi Purhonen, Hans Hebert, Robert T Sauer, Tania A Baker, Ute Römling

## Abstract

Pandemic *Pseudomonas aeruginosa* clone C strains encode a xenolog of FtsH (PaFtsH2), an inner-membrane associated ATP-dependent protease. *FtsH1* supports growth and intrinsic antibiotic resistance but cannot be replaced by *ftsH2*. We show that purified PaFtsH2 degrades fewer substrates than PaFtsH1. Swapping residues of a short MC peptide that links transmembrane helix-2 with the cytosolic AAA+ ATPase module from PaFtsH1 into PaFtsH2 improves hybrid-enzyme substrate processing *in vitro* and enables PaFtsH2 to substitute for PaFtsH1 *in vivo*. FtsH1 MC peptides are glycine rich. Introducing three glycines into the membrane-proximal end of PaFtsH2’s MC linker is sufficient to elevate activity *in vitro* and *in vivo*. Electron microscopy including PaFtsH2 indicates that MC linker identity influences FtsH flexibility. Our findings establish that the efficiency of substrate processing by two PaFtsH isoforms depends on how they are attached to the membrane and suggest that greater linker flexibility/length allows FtsH to degrade a wider spectrum of substrates. As FtsH2 homologs occur across bacterial phyla, we hypothesize that FtsH2 is not a latent enzyme, rather recognizes specific substrates or is activated in specific contexts or biological niches. We hypothesize that such linkers might play a more determinative role in functionality and physiological impact of FtsH proteases than previously thought.

## Introduction

FtsH, a membrane-bound protease and member of the AAA+ (<u>A</u>TPases <u>a</u>ssociated with diverse cellular <u>a</u>ctivities) superfamily of ATPases, is widespread in prokaryotes and in eukaryotic organelles (1–3). Although some bacteria encode five distinct classes of AAA+ proteases, FtsH is typically the only protease that is biologically essential, is the most phylogenetically conserved, and is the only inner-membrane-associated AAA+ protease in most Gram-negative bacteria (4). FtsH is an essential modulator of lipid metabolism in *Escherichia coli* and is important for growth, virulence, production of secondary metabolites, biofilm formation and formation of persister cells in *Pseudomonas aeruginosa* as well as many other bacteria (5–10). Bacteria, including *P. aeruginosa,* typically encode just one FtsH enzyme, but the pandemic clone C group of *P. aeruginosa* is an example of organisms that have undergone ‘*ftsH*-gene expansion’ as it carries a second FtsH variant introduced by horizontal transfer (5, 11).

Like other AAA+ proteases, FtsH functions as a hexamer with an axial channel that serves as a conduit for polypeptide translocation into a degradation chamber. A prototypical FtsH subunit begins with a short N-terminal cytoplasmic segment, followed by a transmembrane helix (TM1), a periplasmic domain, another transmembrane helix (TM2), a cytoplasmic AAA+ substrate unfolding/translocation module, and a cytoplasmic M41-family peptidase domain (Figures 1A and S1). A cytosolic segment of ∼25-residues connects TM2 and the AAA+ module and hereafter is called the MC or membrane-cytosolic linker. As FtsH is always membrane-tethered, it has been suggested that this linker may help gate substrate entry (3). All of these regions of FtsH are required for its function (12–18).

**Figure 1.**
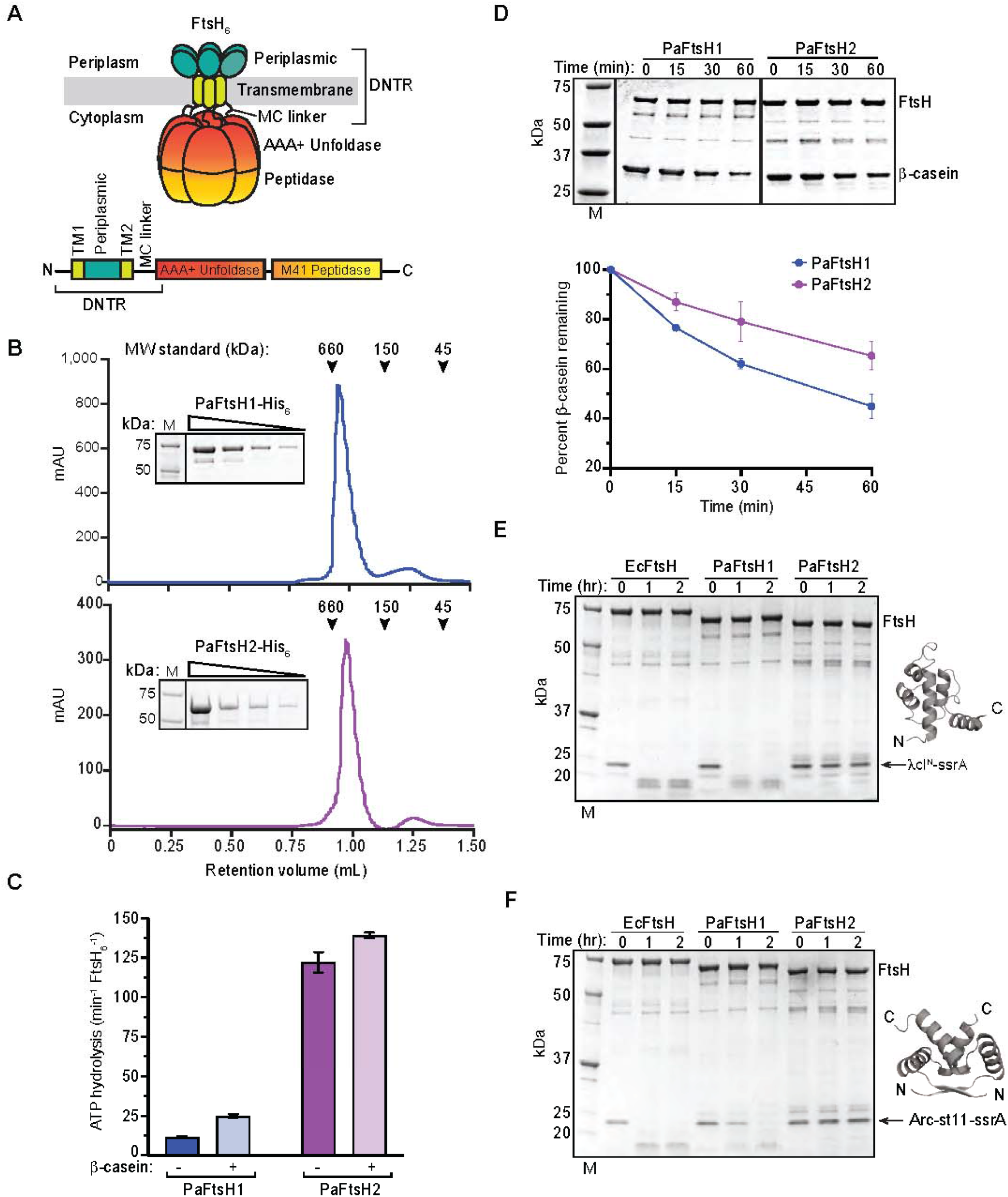
Purified PaFtsH1 and PaFtsH2 have distinct biochemical activities in *vitro*. **A** Schematic (top) and linear representation (bottom) of a prototypical FtsH_6_ enzyme in a bacterial cell with key regions labeled. TM1: Transmembrane helix 1; TM2: transmembrane helix 2; MC: membrane-cytoplasmic; DNTR: diverse N-tenninal region. The DNTR spans the N-terminus. TM1, periplasmic domain. TM2, MC linker, and the first seven amino acids in the AAA+ ATPase domain. **B** Empirical molecular weight determination of PaFtsH1_6_ and PaFtsH2_6_ purified from *E.coli.* Purified PaFtsH1_6_ and PaFtsH2_6_ were run on an analytical Superdex 200 3.2/3.0 gel filtration column and separated bySDS-PAGE in a two-fold dilution series (insets). Molecular weights of PaFtsH enzymes were consistent with hexamer formation plus ∼3.5 NP-40 micelles per PaFtsH1-6xHis hexamer (PaFtsH1-6xHis: 743,373 Da; one NP-40 micelle:∼90 Da) and ∼2.7 NP-40 micelles perPaFtsH2-6xHis hexamer (PaFtsH2-0xH is: 659,792 Da). Theoretical molecular weights of6xH is-tagged PaFtsH monomers: PaFtsH1-6xHis: 70,886 Da; PaFtsH2-6xHis: 69,521 Da. MW: Molecular weight; M: Molecular weight marker. mAU: milli-Absorbance Untt. **C** Hydrolysis of ATP (5 mM) by PaFtsH1_6_ or PaFtsH2_6_ (0.43 µM) in the presence or absence of the substrate β-casein (40 µM) at 40°C. **D** β-casein (40 µM) was incubated at 40°C with PaFtsH1_6_ or PaFtsH2_6_ (0.43 µM) and degradation kinetics were monitored bySDS-PAGE. Reactions contained 5 mMATP and a regeneration system. Data points are averages of three independent replicates± SD (bottom). M: Molecular weight marker. **E** λcI^N^-ssrA(15 µM) was incubated at 40°C wtth PaFtsH1_6_ or PaFtsH2_6_ (3.04 µM) in the presence of5 mMATP and a regeneration system. λcI^N^ PDB: 1LMB (Beamer and Pabo 1992). M: Molecular weight marker. **F** Degron-taggedArc repressor (Arc-st11-ssrA; 15 µM) was incubated at 40°C with PaFtsH1_6_ and PaFtsH2_6_ (3.53 µM) in the presence of 5 mMATP and a regeneration system. Arc-st11 dimer PDB: 1ARR (Bonvin et al, 1994). M: Molecular weight marker.

FtsH recognizes protein substrates by binding to N- or C-terminal degrons and less commonly to internal disordered regions (12, 19, 20). One group of FtsH substrates includes incomplete proteins tagged with a C-terminal ssrA degron, a short peptide sequence attached to nascent chains via tmRNA-mediated ribosome rescue (19, 21, 22). FtsH participates in membrane-protein quality control by degrading misassembled and misfolded inner-membrane proteins and regulates proteotoxic stress by controlling intracellular levels of the RpoH/σ^32^ heat-shock transcription factor (5, 23–27);). In some cases, adaptor proteins influence the degradation efficiency and specificity of FtsH proteases. For example, the heat shock transcription factor RpoH requires the co-chaperone DnaK and the signal recognition particle (SRP) to be efficiently processed ((28–30)). On the other hand, the periplasmic membrane proteins, HflK and HflC, interact with FtsH (31) and negatively regulate proteolysis of SecY (26) and λ cII (32). Other adaptor proteins influence degradation of LpxC and processing of Colicin D by FtsH (8, 33).

As all clone C strains, the virulent aquatic isolate of *P. aeruginosa* SG17M carries a transmissible locus of stress tolerance (tLST) important for survival under multiple stress conditions that encodes a xenolog of FtsH called FtsH2 ((Figure S2A); (11, 34)). We previously found that inactivation of the genomic FtsH copy, FtsH1, causes a substantial growth deficiency and other phenotypes, whereas FtsH2 appears to act primarily as a backup enzyme under the growth and stress conditions tested (5, 35). Here, we investigate the molecular basis of the differential activities and impacts of PaFtsH1 and PaFtsH2. Using phenotypic and biochemical assays, we demonstrate that the N-terminal region of the PaFtsH1 and PaFtsH2 homologs, which include TM1, TM2, the periplasmic domain, and the MC linker, is necessary and sufficient to determine function of the two distinct FtsH enzymes *in vivo*. Even more, inserting just three glycines into the PaFtsH2 linker to mimic the glycine-rich feature of the PaFtsH1 sequence results in a ‘gain-of-function’ phenotype. Notably, purified PaFtsH2 hybrids carrying PaFtsH1-like MC linker alterations gain proteolytic activity against model substrates *in vitro*. We propose a model that explains how the composition, length, and flexibility of the MC linker could modulate FtsH enzyme function by affecting the position and mobility of the cytoplasmic AAA+ protease with respect to the plane of the membrane and control the accessibility of specific substrates to the AAA+ axial channel during the initiation of efficient proteolysis.

## Results

### The N-terminal regions of FtsH1 and FtsH2 are highly divergent

To identify the molecular mechanism of the differential activity of PaFtsH2 compared to PaFtsH1, we first assessed the sequence conservation of the individual domains. The overall sequence identity between PaFtsH1 and PaFtsH2 is 45%, with the most conserved regions being the AAA+ module and protease domain (54% identity). Signature motifs for substrate processing, the pore motif FVG, and catalytic activity are completely conserved in PaFtsH2 and PaFtsH1 for both domains and therefore suggests that any functional differences between the enzymes would not be attributed to differences in these motifs (Figures 1A and S1). The N-terminal ∼150 amino acids are most divergent (∼24% identity) (Figures 1A and S1). We refer to this region from the N-terminus through the first β-strand of the AAA+ module as the divergent N-terminal region (DNTR).

### Purified PaFtsH1 and PaFtsH2 assemble as hexameric ATPases and proteases

To compare the intrinsic biochemical activities of PaFtsH1 and PaFtsH2, we overexpressed both enzymes in *E. coli* and purified them to >95% homogeneity following extraction from cell membranes. Each enzyme eluted from a size-exclusion column as a single peak, with a molecular weight expected for a hexamer bound to detergent micelles (Figure 1B). Both enzymes were active ATPases, with ATP hydrolysis by PaFtsH2 being ∼nine-fold faster than by PaFtsH1 under standard conditions (Figures 1C and S2B). despite being the ‘silent’ gene product *in vivo* and in most *in vitro* instances ((5); and as described below). Addition of casein, a commonly used protease substrate with a molten-globule structure, stimulated the PaFtsH1 ATPase activity about two-fold, but only marginally stimulated that of PaFtsH2 (Figure 1C). Both enzymes effectively degraded β-casein with PaFtsH1 degradation rate being slightly (∼two-fold) faster than PaFtsH2 (Figure 1D). Thus, purified PaFtsH1 and PaFtsH2 are active as ATPases and proteases *in vitro*.

### PaFtsH2 fails to degrade ssrA-tagged proteins

*E. coli* FtsH degrades protein substrates with C-terminal ssrA tags (22, 36). SsrA is a well-recognized automonous degron that can be added to essentially any protein, rendering it a substrate. In the case of SsrA-tagged peptides subjected to degradation by FtsH protease, additional co-factors that aid processing do not seem to be required. Thus, we tested PaFtsH1 and PaFtsH2 degradation of λcI^N^-ssrA, a variant of the N-terminal domain of phage λ repressor (λcI^N^) with the *E. coli* ssrA tag (AANDENYALAA) (37). As assayed by SDS-PAGE, PaFtsH1 degraded this substrate over the course of ∼one hour, whereas PaFtsH2 was essentially inactive in degradation of this substrate (Figure. 1E). Notably, PaFtsH1 did not degrade λcI^N^ lacking the ssrA degron (Figure S3A). PaFtsH1, but not PaFtsH2 also degraded Arc-st11 containing a *P. aeruginosa* ssrA tag (AAND<u>D</u>NYALAA; Figure 1F). Note that throughout this work we found that PaFtsH1 efficiently recognized both the *P. aeruginosa* sequence*-*specific ssrA-tag (-AAND<u>D</u>NYALAA) and the *E. coli* ssrA sequence (-AAND<u>E</u>NYALAA), which differ by just one amino acid.

### PaFtsH1 and PaFtsH2 have similar peptide-binding profiles

As degrons are often unstructured peptides exposed on a protein surface, measuring binding of a AAA+ protease to immobilized peptide arrays can be a useful method for identifying and characterizing degron sequences. However, these binding experiments do have to be followed by functional assays to establish the role of identified sequences in protease recognition of the intact protein. To rapidly investigate if PaFtsH1 and PaFtsH2 had similar or distinct peptide binding properties, we synthesized arrays of 12-residue peptides from known or probable FtsH substrates from *P. aeruginosa*. Known PaFtsH1 substrates include RpoH and PhzC, and PaFtsH1 or PaFtsH2 candidate substrates include GlmM, LepB and MinD; (5, 27). Subsequently, peptide arrays were probed them with ^35S^PaFtsH1 and ^35S^PaFtsH2 (Figs. 2A, 2B, S4, Table S1). Radiolabeled PaFtsH1 and PaFtsH2 bound these arrays with very similar profiles, indicating that both enzymes interact with the immobilized peptides in the same manner. Note that this approach revealed strong binding of *E. coli* FtsH, and both *P. aeruginosa* enzymes to a sequence in RpoH (region 3, Figures 2A, 2B) that is known to be important for degradation in *E. coli* (38, 39) indicating that at least some of the protein-protein interactions identified here are involved in substrate recognition by these enzymes.

**Figure 2:**
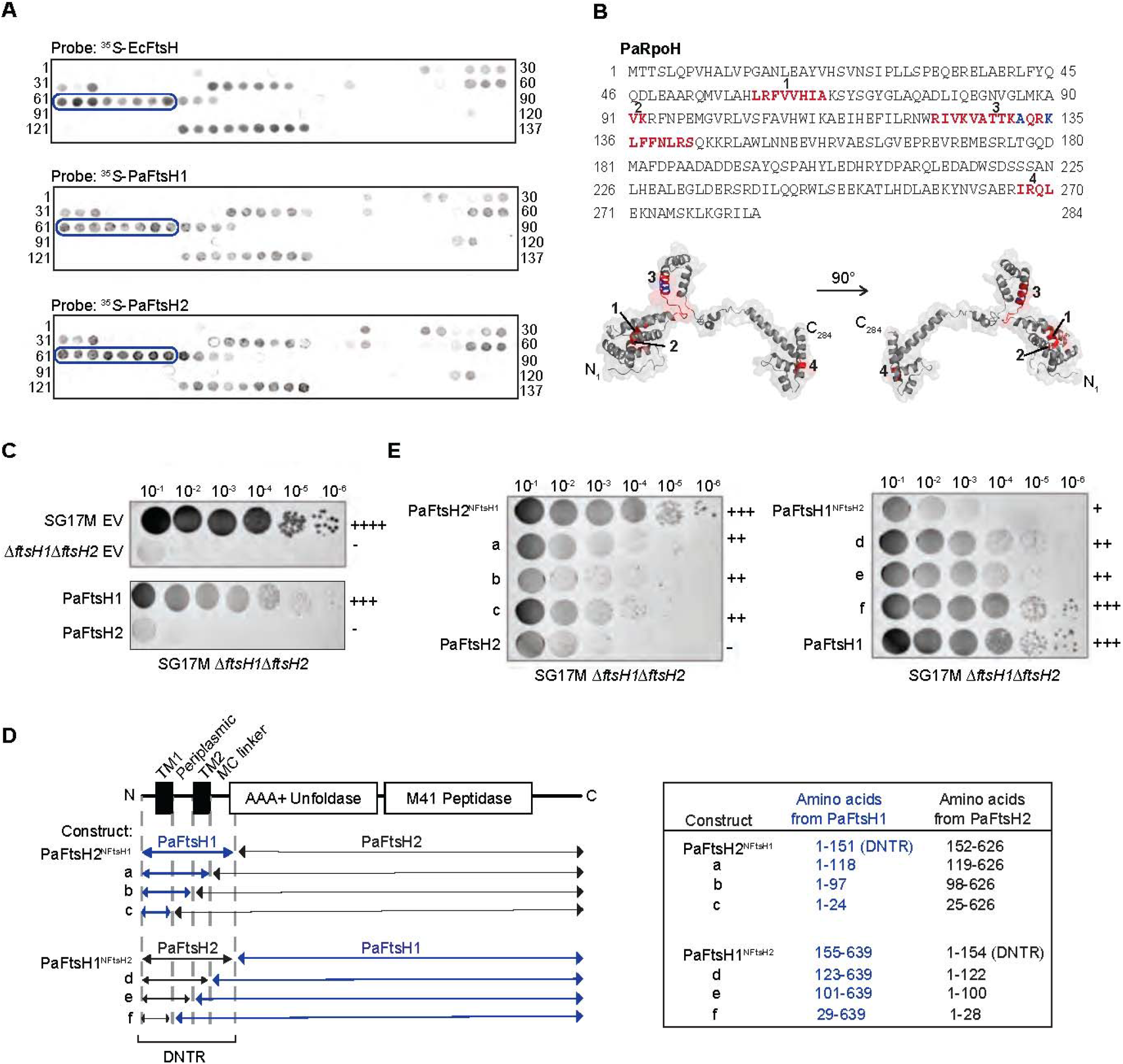
PaFtsH1_6_, and PaFtsH2_6_, have similar *in vitro* peptide binding profiles and differentially affect *in vivo* colony size. **A** Peptide array of the RpoH sequence from *P aeruginosa* (P42378IRpoH_PSEAE: 1-284, sliding window of 12 amino acids with a step size of two amino acids towards the C-terminus with each spot) probed with 1 µM ^35^S-EcFtsH, (top), ^35^S-PaFtsH1, (middle), or ^35^S-PaFtsH2, (bottom) in the presence of 1.25 mM ATPyS. Peptide sequences corresponding to each spot are listed in Table S1. Circled spots indicate peptides containing residues A132 and/or K135, which were previously shown to be important for *in vivo E. coli* FtsH (EcFtsH) binding to RpoH (conserved RpoH residues from E.coli areA131 and K134; Obrist et al, 2009). **B** Amino acid sequence of PaRpoH (top). Residues in red indicate regions bound by ^35^S-EcFtsH_6_, ^35^S-PaFtsH1_6_, ^35^S-PaFtsH2_6_,. Residues in blue correspond to A132 and K135 (see panel A legend). FtsH-binding sequences identified by peptide blotting are mapped on the predicted PaRpoH structure (bottom; Raman et al, 2013) and numbered as on the sequence above. **C** Colony sizes of wild type *P aeruginosa* SG17M, the SG17M Δ*ftsH1*Δ*ftsH2* double deletion and SG17M Δ*ftsH1*Δ*ftsH2* complemented with either an empty expression vector (EV) pJN105 or pJN105-derived expression of PaFtsH1 or PaFtsH2. *PaftsH1* and PaftsH2 genes had been cloned into the expression vector pJN105 under the translational regulation of the PaftsH2 Shine Dalgarno (SD) sequence.“+” and “−” scoring system relates the degree of colony growth of a particular strain genotype to that of SG17M EV(++++; wild type colony growth), or SG17M Δ*ftsH1*Δ*ftsH2* (-; very poor colony growth). For (C) and (E), numbers along the top refer to the serial dilution factor for plating cells. **D** PaFtsH1 and PaFtsH2 hybrid protein constructs mapped linearly onto a schematic of their domain structure with boundaries highlighted. Regions corresponding to PaFtsH1 identity are depicted in blue (left). The amino acid sequences of FtsH1 and FtsH2 comprising the hybrid construct are indicated along with their construct abbreviation (right). TM1: Transmembrane helix 1: TM2: Transmembrane helix 2; MC: Membrane-cytoplasmic; DNTR: Diverse N-terminal region. **E** Colony size of the *P aeruginosa* SG17M Δ*ftsH1*Δ*ftsH2* double deletion complemented with Pa*ftsH1* and Pa*ftsH2* hybrid constructs as introduced in panel D. “+”and“-” scoring system is the same as in (C).

### The C-terminal region of the DNTR determines *in vivo* impact

We have reported previously that although both PaFtsH1 and PaFtsH2 are produced in exponential phase (as well as under numerous other growth medium conditions), PaFtsH1 plays an almost exclusive role in supporting robust growth, as measured by colony size and growth rate (5). Figure 2C recalls that *ftsH1* but not *ftsH2* expression rescues the tiny colony size phenotype of SG17M Δ*ftsH1*Δ*ftsH2*. To determine if the DNTR, the most distinct region between these two proteins, determines functionality under these experimental conditions, we expressed constructs containing residues 1-151 of PaFtsH1 (PaFtsH1 DNTR) appended to the “body” of PaFtsH2 (PaFtsH2^NFtsH1^) or residues 1-154 of PaFtsH2 (PaFtsH2 DNTR) attached to “body” of PaFtsH1 (PaFtsH1^NFtsH2^) under translational control of the *ftsH2* Shine-Dalgarno sequence in SG17M Δ*ftsH1*Δ*ftsH2*. Notably, PaFtsH2^NFtsH1^ rescued the colony size defect whereas PaFtsH1^NFtsH2^ did not (Figures 2D and 2E), indicating that the DTNR of *ftsH1* can function with the AAA+ module and M41 peptidase of *ftsH2* to support robust growth. To identify shorter regions of the PaFtsH1 DTNR that contribute most strongly to this functionality, we constructed and tested additional PaFtsH1 and PaFtsH2 chimeras with iteratively swapped DTNR segments (Figures 2D and 2E). We found that restoration of colony size was largely determined by an amino acid stretch of PaFtsH1 that included the MC linker and the N-terminal seven residues of β-strand 1 in the AAA+ module (see especially Figure 2E, constructs a and d). On the other hand, the equivalent amino acid stretch of PaFtsH2 abolished the restoration of colony size by PaFtsH1. We call this segment MC^7+^.

### MC sequences from PaFtsH1 activate PaFtsH2 degradation

In order to unambiguously assess the contribution of the MC^7+^ to functionality, we constructed, expressed, and purified a variant called PaFtsH2^H1-link-32^ containing the 32-residue MC^7+^ region from PaFtsH1 with all other enzyme sequences derived from PaFtsH2 (Figures 3A, 3B. Strikingly, PaFtsH2^H1-link-32^ degraded λcI^N^-ssrA, whereas the parental PaFtsH2 protease did not, and also degraded this substrate at a rate faster than PaFtsH1 in an ATP-dependent and ssrA tag-dependent reaction (Figures 3C, S3A, S3B). However, PaFtsH2^H1-link-32^, like its PaFtsH2 parent, was unable to degrade Arc-st11-ssrA (Figure 3D), suggesting that MC^7+^ region from PaFtsH1 is not directly involved in recognition of the ssrA tag or that such recognition is highly dependent on features of the attached protein. PaFtsH2^H1-link-32^ degradation of λcI^N^-ssrA generated a novel ∼5.7 kDa product (marked by asterisks in Figures 3C, 3F, S3A, S3B) that mass spectrometry identified as residues 1-51 of λcI^N^-ssrA, as expected if proteolysis initiates from the C-terminal ssrA degron of this substrate (Figure S3C). The presence of this degradation product could represent a difference in peptide-cleavage specificity within the protease chambers of PaFtsH1 and PaFtsH2 or a difference in substrate processing by the AAA+ modules of these enzymes. Indeed it has been observed that FtsH can perform limited proteolytic cleavage (33).

**Figure 3:**
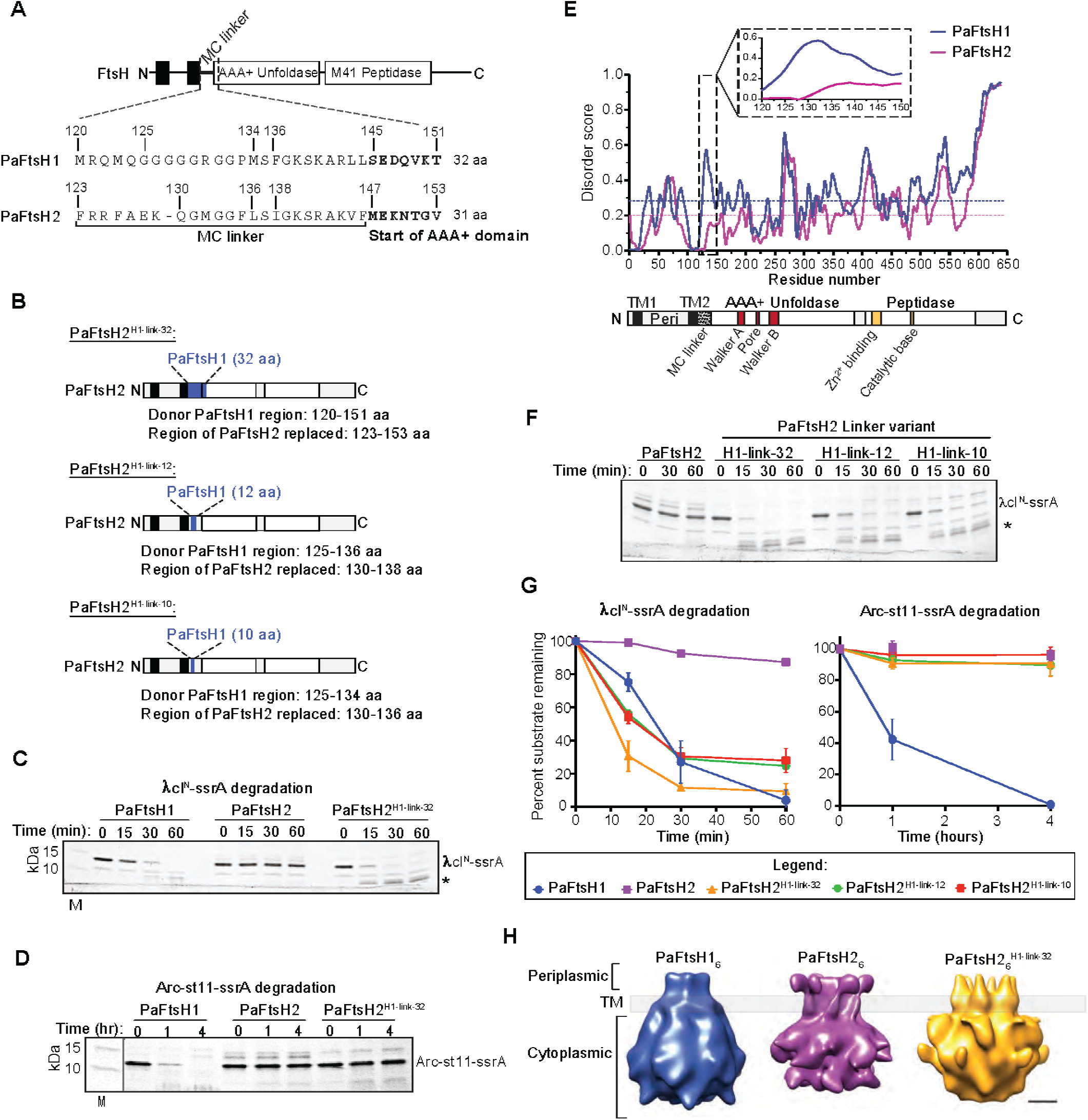
The membrane-cytoplasmic (MC) linker of PaFtsH1_6_ activates degradation by PaFtsH2_6_ *in vitro*. **A** Linear map of a model FtsH enzyme with a zoom-in on MC linker sequences from PaFtsH1 (32 amino acids) and PaFtsH2 (31 amino acids). Bolded residues denote the start (N-terminal side) of respective PaFtsH1 and PaFtsH2 AAA+ATPase domains. **B** Schematic of FtsH2 MC linker chimera constructs: PaFtsH2^H1-link-32^, PaFtsH2 ^H1-link-12^, PaFtsH2 ^H1-link-10^. Regions in blue represent residues of PaFtsH1 identity on an otherwise PaFtsH2 body. The number of amino-acids derived from PaFtsH1 are indicated in the superscript of the variant name. Black rectangles represent TM domains, and white rectangles indicate the AAA+ unfoldase and M+1 peptidase regions, respectively. **C** Degradation of λcI^N^-ssrA(15 μM) by PaFtsH1_6_, PaFtsH2_6_, and PaFtsH2_6_ ^H1-link-32^ (3.0+ pM) as described in Figure 1E. Bands corresponding to full-length λcI^N^-ssrA substrate and a ∼5.7 kDa degradation intermediate product (denoted by asterisk) are indicated. **D** Degradation of Arc-st 11-ssrA(15 pM) was monitored in the presence ofPaFtsH1_6_, PaFtsH2_6_and PaFtsH2_6_ ^H1-link-32^ (3.53 pM) as described in Figure 1F. **E** Structural disorder prediction profiles of PaFtsH1 (blue) and PaFtsH2 (purple) as predicted bythe lUPred algorithm (Dosztanyi et al, 2005). Dotted lines indicate the average disorder scores for residues 1-590 of PaFtsH1 (blue, 0.29) and PaFtsH2 (purple, 0.20). Inset shows a zoom-in ofthe MC^7+^ linker regions. Bottom schematic is a linear model of an FtsH enzyme with important structural and functional regions noted on the diagram. Abbreviations are as follows: TM1: Transmembrane domain 1; Peri: Periplasmic domain; TM2: Transmembrane domain 2. **F** Degradation of λcI^N^-ssrA(15 pM)was monitored in the presence ofPaFtsH1_6_, PaFtsH2_6_ and PaFtsH2_6_^H1-link-32^ (3.0 + pM) as described in Figure 1E. **G** Quantification of full-length substrate remaining over reaction time courses. Error bars represent SD of experiments performed in triplicate. Representative degradation kinetics as monitored bySDS-PAGE by PaFtsH1_6_ and PaFtsH2_6_ are shown in Figures 1E and 1F. **H** 3D-reconstruction from negative stain TEM of PaFtsH1_6_, PaFtsH2_6_, and PaFtsH2_6_^H1-link-32^. Scale bar: 3 nm. Periplasmic, transmembrane (TM), and cytoplasmic domains are noted on figure.

We continued this protein dissection approach and constructed and purified enzymes containing most of PaFtsH2 and just 10 or 12 residues of the MC linker of PaFtsH1 (Figure 3B). Importantly, both chimeras degraded λcI^N^-ssrA like PaFtsH2^H1-link-32^, but unlike the parental PaFtsH2 (Figures 3F, 3G, S3A). The MC linker of PaFtsH1 is predicted by computational methods to be more disordered/flexible than that of PaFtsH2, as are the linker segments containing just 10 or 12 PaFtsH1 residues (Figs. 3E, S3D, and S5).

### The MC linker appears to contribute to conformational flexibility

To investigate whether the functional differences we observed with the different MC linkers was reflected in differences in enzyme structure/dynamics, we imaged detergent-solubilized PaFtsH1, PaFtsH2, and the most active hybrid protein PaFtsH2^H1-link-32^ by electron microscopy and negative staining, resulting in fields of particles representing single hexamers (Figure S6A). Reconstructed 3D-maps using six-fold symmetry produced distinct ∼22-Å resolution density maps for the three enzymes (Figure 3H). Each map contained obvious density for the large cytoplasmic domains and weaker density for the periplasmic domain, whereas the transmembrane helices were obscured by detergent and not well-visualized (Figure S6B).

Although PaFtsH1 and PaFtsH2 showed cone-shaped appearances, more defined-shape features were visible for PaFtsH2, including well-defined domain-like shapes in the periplasmic ring and clear delineation between the AAA+ ATPase and peptidase segments of the cytoplasmic regions (Figure 3H). We interpret these structural differences as an indication of PaFtsH2’s relative conformational homogeneity, whereas PaFtsH1 may adopt more conformations and thus “blur” structural features. These observations are consistent with the hypothesis that the shorter, stiffer MC linker in PaFtsH2 confines its movements, potentially hampering substrate engagement and/or unfolding. Interestingly, the structural definition in the PaFtsH2^H1-link-32^ variant was intermediate between those of PaFtsH1 and PaFtsH2. Especially noticeable in PaFtsH2^H1-link-32^ was the loss of cytoplasmic-ring definition compared to parental PaFtsH2. These results also support the notion that the PaFtsH1 MC linker imparts greater ability to “explore” more conformations than the PaFtsH2 linker.

The PaFtsH2 structure was more similar to a recent cryo-EM structure of *Aquifex aeolicus* FtsH ((3); EMD-11161) than the other structures, probably because it had the most well-defined periplasmic and cytoplasmic domains. Of note, AaFtsH is predicted to possess a more rigid linker than PaFtsH1 with three glycine residues. The crystal structure of the *A. aeolicus* cytoplasmic domain of FtsH (PDB 4WW0; (40)) also fit best with PaFtsH2, although the AAA+ modules and peptidase domains could be acceptably fit in each of the three density maps (Figure S6B).

### FtsH1 MC linker length determines complementation strength by H1-H2 chimeras

PaFtsH2^H1-link-32^ complementation of SG17M Δ*ftsH1*Δ*ftsH2* colony size was less complete than PaFtsH2^NFtsH1^ complementation but far greater than complementation by PaFtsH2 alone (Figure 4A). PaFtsH2^H1-link-10^ did not rescue colony growth. We also monitored the ability of different FtsH linker chimeras to complement tobramycin susceptibility and growth in liquid M63-citrate minimal media (Figures 4B-D). Expression of PaFtsH2^H1-link-32^ but not PaFtsH2^H1-link-10^ restored tobramycin tolerance to a level similar to that afforded by PaFtsH1 or PaFtsH2^NFtsH1^ (Figure 4B). Similarly, only PaFtsH2^NFtsH1^ and PaFtsH2^H1-link-32^ partially complemented the slow-growth defect of SG17M Δ*ftsH1*Δ*ftsH2* in M63-citrate medium better than PaFtsH2 itself (Figures 4C and 4D). Thus, in *vitro* and *in vivo* activities of hybrid proteins are not entirely congruent, as distinct substrate(s) or protein conformation might be determinative for recovery of *in vivo* phenotypes such as colony size.

**Figure 4:**
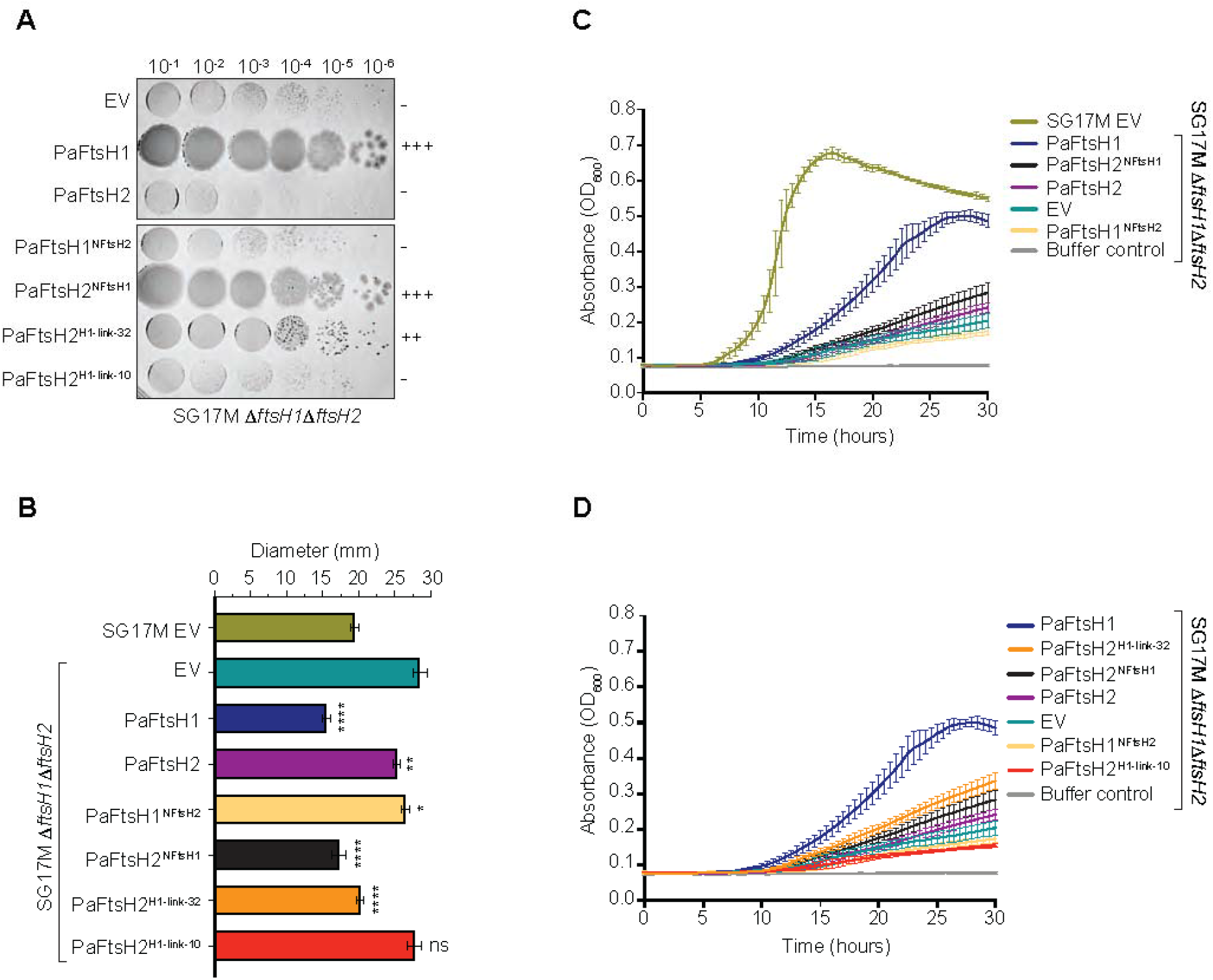
MC linker identity determines the strength of complementation in *P. aeruginosa* SG17M Δ*ftsH1*Δ*ftsH2* cells. **A** Colony growth of P. aeruginosa SG17M AftsH1AftsH2 double deletion strains upon complementation with an empty expression vector control pJN105 (EV) PaFtsH1, PaFtsH2, PaFtsH1^NFtH2^, PaFtsH2^NFtH1^, PaFtsH2^H1-link-32^ or PaFtsH2^NFtH1^ as indicated. Strains were grown overnight at 37°C on LB-agar plates with 30 pg/mL gentamicin.“+” and scoring system relates the degree of colony growth of a particular strain genotype to that of SG17M Δ*ftsH1*Δ*ftsH2* complemented with PaFtsH1 (+++) or PaFtsH2 (-). **B** Measurement of zone of inhibition upon treatment with the aminoglycoside antibiotic tobramycin. *P. aeruginosa* SG17M *&ftsH1&ftsH2* double deletion strains were complemented by expression with PaFtsH1 and PaFtsH2 wild type and hybrid constructs as indicated. Wild type SG17M strain harboring an empty expression vector (EV) is shown for reference. Abacterial suspension of 0.5 Farland was distributed onto a Mueller Hinton agar plate containing 30 pg/mL gentamicin. The diameter of the zone of growth Inhibition was measured after Incubation at 37°C for 20-24 hours in the presence of a 10 pg tobramycin disk. ****: p < 0.00002 ***: p < 0.0002: **: p < 0.002: * p < 0.02 (compared to “EV” (teal bar)). **C-D** Growth of *P aeruginosa* SG17M wild type or Δ*ftsH1*Δ*ftsH2* double deletion strains complemented by plasmid-borne expression of the indicated PaFtsH variants in M63 citrate minimal media. EV: pJN105 empty expression vector control.

### MC linker evolution

Previous phylogenetic analyses of the closest homologs of PaFtsH1 and PaFtsH2 in different taxonomic groups of bacteria reveled that similarity among FtsH1 genes/proteins was largely in line with overall phylogenetic relationships (compared to 16S rRNA), whereas FtsH2 similarity was not. These data are congruent with *ftsH2* genes horizontally transferred as part of the tLST with an apparent bias toward pathogens into distinct genetic backgrounds of bacterial species including *P. aeruginosa*. *Escherichia coli* and *Klebsiella pneumoniae* (5, 11, 34).

Given our structure/function data on the impact of the MC linker in PaFtsH1 versus PaFtsH2 activity, we hypothesized that MC linker similarity is congruent with FtsH protein clusters. To this end, we aligned representative FtsH proteins with similarity to PaFtsH1 and PaFtsH2 from the NCBI and UniProt databases and additionally included selected FtsH representatives from established major phyla of the bacterial phylogenetic tree. As FtsH proteins are highly conserved also in other major evolutionary branches including amoeba, fungi, plants and mammals, FtsH representatives from select organisms, such as the model plant *Arabidopsis thaliana* (which possess 11 FtsH homologs) and *Homo sapiens*, were included in our analysis. We generated a protein evolution tree where branch length represents the number of sequence changes between entries (Figure 5A; (5)). As expected, the FtsH1 and FtsH2 representatives each clustered together, but far apart from each other. FtsH1 proteins (blue branches) and FtsH2 proteins (purple branches) were therefore used to generate two sequence alignments, to allow comparison between each family’s MC linkers.

**Figure 5:**
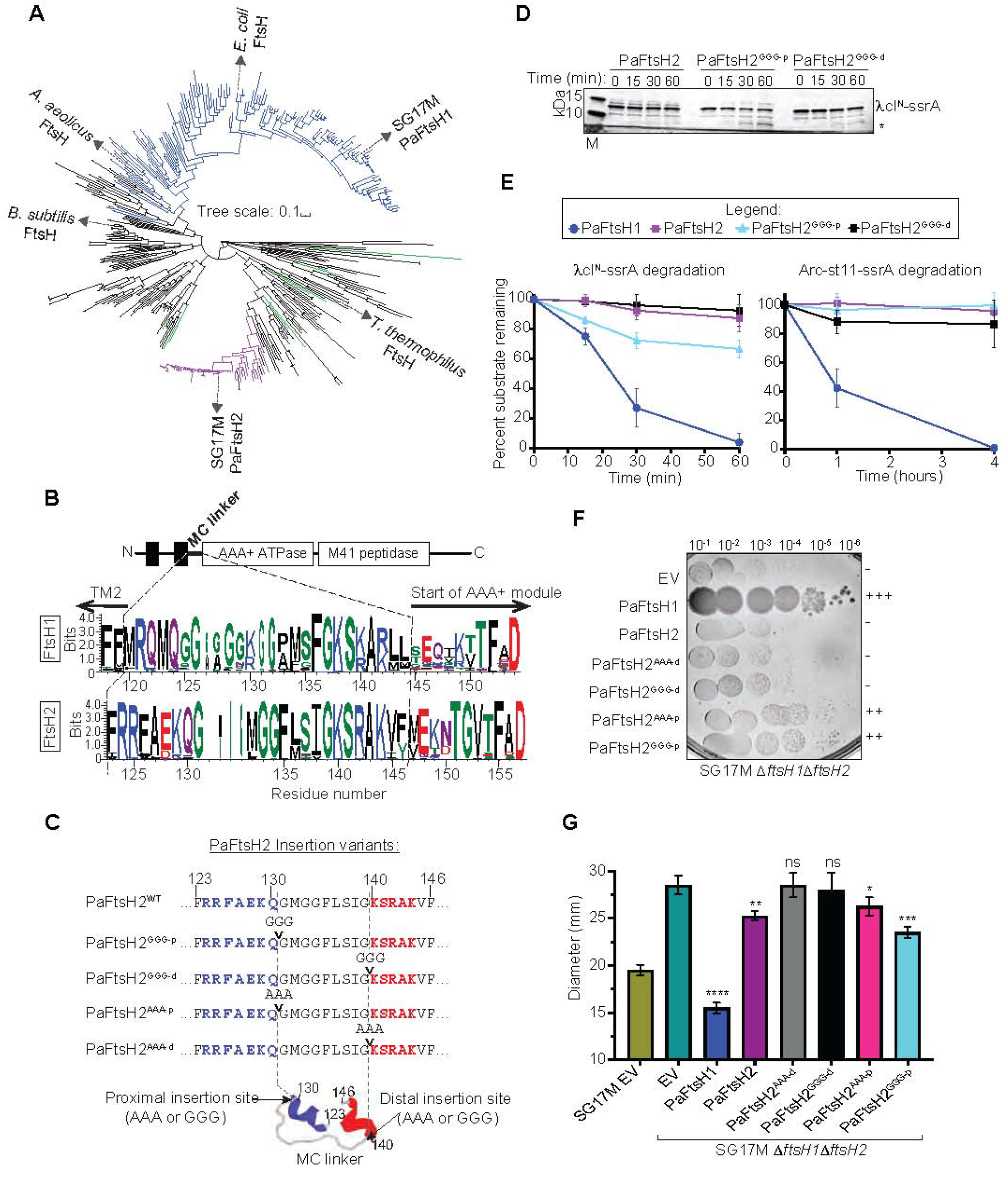
Insertion of triplicate alanine or glycine residues in the proximal region of the PaFtsH2 MC linker activates PaFtsH2 *in vivo* and *in vitro*. **A** Evolutionary tree of full length FtsH proteins from each bacterial phylum and representative FtsH proteins spanning archaeal and eukaryotic kingdoms (see Table S2) FlsH proteins from select species are labeled. Blue branches indicate FtsH proteins with a FtsH 1-like linker. purple branches indicate FtsH proteins with a FtsH 2-like linker and green branches indicate *Arab1dopsis thaliana* FtsH proteins Tree scale indicates amino acid substitutions per site **B** Domain structure of a model FtsH enzyme and WebLogos of FtsH1 and FtsH2 MC linker sequences constructed from 149 sequences ofFtsH1 (blue branches in A)) and 48 sequences of FtsH2 (purple branches in A)) **C** Domain structure of PaFtsH2 with a zoom-in on the MC linker sequences of indicated PaFtsH2 triplicate residue insertion vanants PaFtsH2 MC linker structural prediction was generated by PEP-FOLD 3.5 (bottom, Lamiable et al. 2016)where blue and red helices indicate predicted proximal and distal helices, respectively, and the course grained energy (sOPEP) is -35 74 Site of proximal insertion is between amino acids 131-132, and site of distal insertion is between amino acids 139-140. ArTows indicate positions of residue insertions (Gly-Gly-Gly (GGG) or Ala-Ala-Ala (AAA)) for four PaFtsH2 insertion variants PaFtsH2^GGG-p^, PaFtsH2^GGG-d^, PaFtsH2^AAA-p^, PaFtsH2^AAA-d^ **D** Degradation of λcI^N^-ssrA (15 µM) was monitored in the presence of PaFtsH2, PaFtsH2 ^GGG-p^, or PaFtsH2 ^GGG-d^ (3.04 µM hexamer equivalents) as described in Figures 1E and 3C. Bands corresponding to ful length λcI^N^-ssrA and a ∼5.7 kDa degradation intermediate product (denoted by asterick) are indicated. M Molecular weight marker. **E** Ouantification of full-length substrate (λcI^N^-ssrA and Arc-st11-ssrA) remaining over reaction time courses. ErTor bars represent SD of expenments performed in triplicate Data for degradation reactions containing PaFtsH1_6_ and PaFtsH2_6_ are the same as in Figure 3G. **F** Colony growth of *P aerugmosa* SG17M Δ*ftsH1*Δ*ftsH2* double deletion strains complemented by expression of the indicated PaFtsH variants. **G** Measurement of zone of inhibition upon treatment with the aminoglycoside antibiotic tobramycin *P aeruginosa* SG 17M Δ*ftsH1*Δ*ftsH2* double deletion strains were complemented by expression with PaFtsH1 and PaFtsH2 wild type and triplicate insertion constructs as indicated Wild type SG17M strain harboring an empty expression vector [EV) is shown for reference ****: p < 0.00002; ***: p < 0.0002; **: p < 0.002; *p < 0.02 (compared to “EV” (dark teal bar))

Comparing the amino acid preferences within the MC linker sequences (including the first seven residues of the AAA+ modules) shows a strikingly-conserved glycine-rich character of the PaFtsH1-type linker in contrast to the PaFtsH2-type linker (Figure 5B). Whereas up to eight clustered glycines (seven being common, *e.g. GGGGG[K/R]GG*) occurred in FtsH1-type linkers, those of the FtsH2 proteins contained fewer glycines and no long Gly-rich blocks (*e.g. GMGG* being common). In addition, the Gly-rich sequence tracks in both types of linkers are positioned differently with respect to the TM2 transmembrane helix and AAA+ modules which also extends the length of TM2 in PaFtsH2 (Figure 5B). However, the FtsH linker has been subject to evolution already in isolates from deeply branching phyla such as Bipolaricaulota and Caldiserica (Figure S7). Although most isolates display FtsH proteins with short glycine-poor linker sequences, long glycine-rich linkers occasionally occur.

### Insertion of Gly-Gly-Gly proximal to the MC linker activates PaFtsH2

The MC linker in PaFtsH2^H1-link-10^ contains more glycines than that of wild-type PaFtsH2, is longer by one residue, and activates proteolysis of λcI^N^-ssrA (Figures 3F, 3G). To probe linker length, flexibility, and positional effects of a 3xglycine stretch within the linker, we cloned, overexpressed, and purified PaFtsH2 variants with three glycines or alanines inserted immediately proximal (PaFtsH2^GGG-p^ and PaFtsH2^AAA-p^) or distal (PaFtsH2^GGG-d^ and PaFtsH2^AAA-d^) to the wild-type MC linker, directly at the sequence junctions between structured and unstructured regions of the predicted PaFtsH1 MC linker peptide fold (Figure 5C). PaFtsH2^GGG-p^, but not PaFtsH2^GGG-d^ partially degraded λcI^N^-ssrA over the course of one hour (Figure 5D, 5E). Consistent with this, plasmid-borne expression of both PaFtsH2^GGG-p^ and PaFtsH2^AAA-p^ partially rescued the colony-size defect of SG17M Δ*ftsH1*Δ*ftsH2*, albeit less well than PaFtsH1, but better than the PaFtsH2 parent (Figure 5F). Similarly, PaFtsH2^GGG-p^ performed better than PaFtsH2^AAA-p^ in the tobramycin assay (Figure 5G). Neither PaFtsH2^GGG-d^ or PaFtsH2^AAA-d^ rescued colony growth (Figure 5F), scored well in the tobramycin assay (Figure 5G), or complemented growth in M63-citrate liquid medium (Figure S8A). However, both linker variants had reduced rates of ATP hydrolysis which might account for their poor activities (Figure S8B). In combination, these results strongly suggest that MC linker length and flexibility influence FtsH biological activity in ways that depend on the position of a flexible segment within the linker.

The fact that PaFtsH2^GGG-p^ has biological activities that PaFtsH2 lacks suggests that the wild-type MC linker of PaFtsH2 lacks flexibility (Figure 6), but this model fails to explain why PaFtsH2^H1-link-10^ was less active than PaFtsH2^GGG-p^ in some *in vivo* assays. We noticed, however, that a glutamine at position 130 in PaFtsH2, near the proximal end of the MC linker (position 124 in PaFtsH1), is highly conserved in the sequence alignments of FtsH2, although not necessarily in all FtsH linkers distinct from FtsH1 (Figures 5B, S1 and S7). As a glutamine at this position is missing in PaFtsH2^H1-link-10^ (Figure 3B), we constructed an additional chimera with five residues from PaFtsH1 that preserves the glutamine at PaFtsH2 position 130 (PaFtsH2^H1-link-5^; Figure 6A). PaFtsH2^H1-link-5^ rescued the colony growth defect as well as PaFtsH2^H1-link-32^ (which also retains Gln^130^) (Figure 6B). As structural predictions place Gln^130^ at the base of the TM2 helix (Figure 6C), it is possible that Gln helps with establishing a structural boundary between the membrane and MC linker, thereby altering enzyme flexibility.

**Figure 6:**
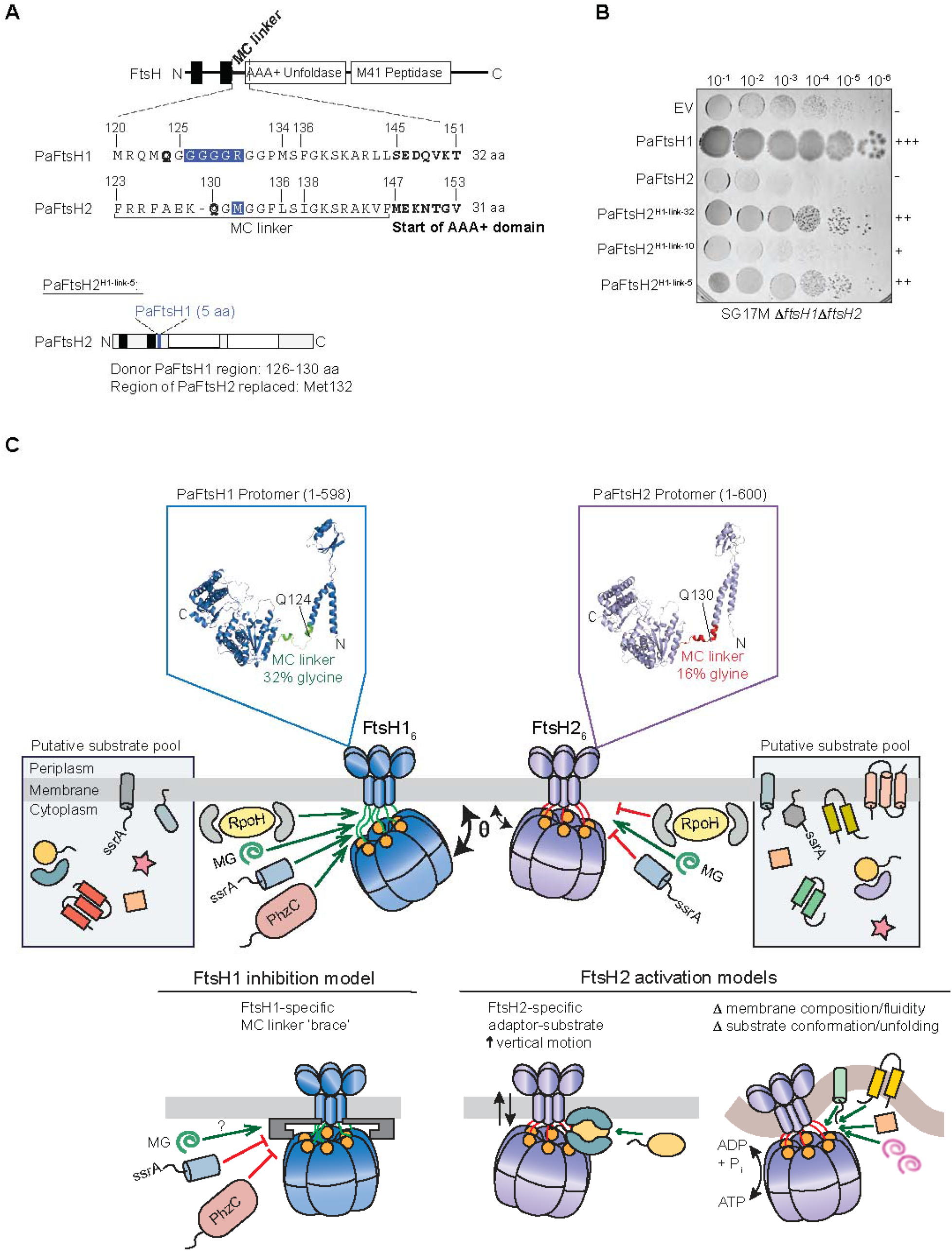
Membrane-cytoplasmic linker properties are important in determining FtsH1 and FtsH2 functionality. **A** Schematic description of the PaFtsH2^H1-link-5^ variant. Mel132 of PaFtsH2 (blue highlight) was replaced with a stretch of five amino acids from PaFtsH1. GGGGR (residues 126-130, blue highlight). PaFtsH1 0124 and PaFtsH2 0130 are denoted in bold and are predicted to localize at the base of TM2 in both enzymes, as referred to in panel C. **B** Colony growth of *P aeruginosa* clone C SG17M Δ*ftsH1*Δ*ftsH2* double deletion strains complemented with either empty vector (EV), wild type PaflsH1, PaflsH2, or the hybrid constructs PaFtsH2 ^H1-link-32^, PaFtsH2 ^H1-link-10^, and PaFtsH2 ^H1-link-5^, as indicated. The plate in this panel is the same as in Figure 4A for EV, PaFtsH1, and PaFtsH2. **C** Model of MC linker-dependent functionality of FtsH proteases. The more flexible MC linker of PaFtsH1 compared to that of PaFtsH2 (32% and 16% glycine, respectively) allows enhanced motion relative to the inner membrane plane () which may promote access of substrates into the FtsH1 axial pore (highly conserved FVG pore loops are indicated by orange-filled circles). The comparatively reduced MC linker flexibility of FtsH2 restricts its plane of motion and inhibits the productive entry of substrates for proteolysis by FtsH2. FtsH2 degrades molten-globule-like (MG) substrates in vitro, which may indicate an in vivo prerequisite for the remodeling of folded substrates prior to FtsH2-dependent degradation. Alternatively, the higher ATPase activity/substrate pulling force of FtsH2 may point to a scenario in which FtsH2 preferentially proteolyzes integral membrane proteins compared to FlsH1, a substrate class that we did not test in vitro. As *ftsH2* is found on a ILST island which confers enhanced tolerance to heat and other stresses to the host organism, ii is possible that a higher membrane fluidity characteristic of higher temperatures is required for the optimal function of FtsH2, or that condition-dependent adaptors or factors are required for the specific delivery of substrates. Although FtsH2 has a more rigid MC linker than that of FtsH1 and its horizontal movement relative to the membrane plane is constrained, it contains a longer TM2 domain which may result in higher vertical ‘up-and-down’ movement within the inner membrane. This vertical movement may be important for the degradation of certain substrates or for regulating substrate entry into the FtsH2 axial pore. As PaFtsH1 was active against the majority of protein substrates tested in our study, models for its inhibition in vivo include those that restrict or “brace” the flexibility of its MC linker such that entry into its axial pore is no longer accessible. Monomer structural predictions of PaFtsH1 (lop left, O9HV48_PSEAE) and PaFtsH2 (top right, A0485GIM3_PSEAI) were generated using AlphaFold. Although the structures are torsionally inaccurate, they are helpful for viewing the proximity of the MC linkers to the transmembrane (towards the N-terminus) and cytoplasmic (towards the C-terminus) domains. Conserved Gin (0) residues as mentioned in Panel A are located at the base of TM2 of each protein and labeled with arrows. The UniProt ID used for modeling PaFtsH2 is 99.52% identical to the PaFtsH2 sequence from SG17M clone C. The three residues that differed from PaFtsH2 (S127, M447, E524) were changed to residues of PaFtsH2 identity using PyMol (S127R, M447I, E524A). Only residues 1-598 of PaFtsH1 and 1-600 of PaFtsH2 are shown in the figures, as the C-terminal regions are predicted to be highly unstructured.

## Discussion

Bacterial species occupy defined ecological habitats based on their physiological and metabolic properties and ability to tolerate different environmental stresses. *P. aeruginosa* is not only a successful environmental species (41), but an important opportunistic human pathogen that infects immune-compromised individuals and contributes substantially to the morbidly and mortality of cystic fibrosis patients (42). Of note, *P. aeruginosa*has been recently found to be one of the most frequently co-infecting bacteria in COVID-19 patients contributing to severity of disease (43, 44). Elucidating the molecular mechanisms that contribute to *P. aeruginosa* clones, groups of closely related strains, with distinct habitat preferences is therefore an important goal. Clone C of *P. aeruginosa* is especially robust and abundant in the environment, causes acute and chronic human infections, and harbours the tLST genomic island, including *ftsH2*, as part of the clone C core genome which provides increased tolerance against elevated temperature and other stresses (11, 41, 45–48). The presence of the FtsH2 xenolog of core genome FtsH distinguishes *P. aeruginosa* clone C from model strains *P. aeruginosa* PAO and PA14 and most investigated model bacterial species, which encode only a single membrane-bound FtsH enzyme. Evolutionary analysis reveals that a very similar genomic islands including an *ftsH2* gene have transferred into and been maintained in clones of other bacterial species including pathogens (11). However, expansion of the FtsH protease family is not uncommon as, e.g. cyanobacteria harbour up to four highly diverse FtsH proteins which cluster with FtsH from other bacterial species (Figure S7). Furthermore, phytoplasma, such as the vineyard-grape pathogen *Flavescence doree*, carry multiple virulence-associated, but clearly paralogous, *ftsH* genes despite having minimalized genomes (Figure S7 (49, 50)). Thus, expansion of FtsH protease variants appears to be an adaptive evolutionary strategy that has occurred more than once and by various mechanisms.

Building upon a previous study (5), here we characterize differences between the FtsH1 and FtsH2 proteases of *P. aeruginosa.* Notably, although both enzymes are active ATP-dependent proteases, we find that they have distinct substrate specificities. For example, both enzymes degrade β-casein, whereas only PaFtsH1 degrades λcI^N^-ssrA and Arc-st11-ssrA. We also demonstrate that many of the biochemical and biological activities of these FtsH paralogs depend on the identity and character of a short region of the linker that connects the TM2 transmembrane helix to the cytoplasmic AAA+ module. While tilting of the catalytic domains against the membrane plane provides substrate access (3), we show in this work that differential MC linker sequences have a much more determinative and flexible role in substrate processing. Future work will identify natural substrate processed by FtsH proteases with distinct linker sequences.

### MC linker flexibility and control of FtsH function

We initially discovered that a hybrid protein containing the TM1 helix, periplasmic domain, TM2 helix, and MC linker region of PaFtsH1 and the AAA+ module and protease domain of PaFtsH2 had cellular activities similar to PaFtsH1 and vice versa. Further sequence swaps identified a short sequence stretch of the cytoplasmic MC linker as being substantially responsible for functionality of the PaFtsH2 catalytic subunits such as FtsH1-like substrate degradation *in vitro* and phenotypic recovery *in vivo*. Indeed, swapping as few as five residues from the PaFtsH1 MC linker into FtsH2 was sufficient to endow the chimeric protease with substantial FtsH1-like activities (Figure 6A, B).

In the hexameric FtsH enzyme, the MC linkers appear to determine the flexibility of the cytoplasmic parts of the protease relative to the membrane (Figure 6B). This model is consistent with ***(i)*** predictions of the flexibilities of different linkers that support or fail to support certain FtsH1-specific functions; ***(ii)*** with the higher glycine content of the MC linker of PaFtsH1 relative to PaFtsH2; ***(ii)*** with low-resolution EM structures of PaFtsH1, PaFtsH2, and an active chimeric enzyme; and ***(iv)*** with the observation that position-specific insertion of just three glycines or alanines into the MC linker of FtsH2 allowed this chimera to partially restore defects caused by an *ftsH1* deletion.

Although our results support the hypothesis that higher MC linker flexibility correlates with FtsH1-like biochemical and biological functions, identification of endogenous biological substrates of *P. aeruginosa* FtsH2 remains to be determined, as does addressing the question of how the properties of the FtsH2 MC linker contribute to its proteolytic specificity? Intracellular levels of *P. aeruginosa* FtsH2 increase during nutrient deprivation and it represents the predominant FtsH species in late-stationary phase (5). Thus, FtsH2-specific substrates may be more prevalent in late-stationary phase and during biofilm formation, possibly as a consequence of its faster rate of ATP hydrolysis compared to PaFtsH1 and/or differences in substrate specificity. ATP-hydrolysis activity by *P. aeruginosa* FtsH2 also has a temperature optimum that is higher than that of the PaFtsH1 enzyme (Figure S2B). *P. aeruginosa* skin infections in humans is a frequent issue associated with hot tub usage and outbreaks on thermally sterilized endoscopes have occurred (51, 52); these high-temperature environments may favor/activate PaFtsH2 due to its more rapid ATPase rate as well as perhaps increased membrane fluidity or change in composition, which could suppress the apparent ‘negative gating’ properties of its ‘rigid’ MC linker and thus contribute to survival of the organism (Figure 6B).

Moreover, the presence of protein adaptors able to alter FtsH1 or FtsH2 activity and function (*e.g.*, HflC and HflK) may also change depending upon specific growth or environmental conditions. An interesting and testable hypothesis is that some adaptors/regulators exert their influence on FtsH activity, at least in part, by abrogating or mimicking the effects of a flexible versus stiff attachment to the inner membrane. Surprisingly, such physiological impact does seem to have developed gradually as selectively investigated deeply branching bacterial phyla contain predominantly, but not exclusively, FtsH proteins with short linkers (Figure S7A). Like the MC linker of *P. aeruginosa* FtsH2, MC linkers from extremophile variants of FtsH (e.g., *A. aeolicus* and *T. thermophilus*) or single-membrane Gram-positive bacteria (e.g., *M. tuberculosis*) also have fewer glycines (Figure S1). This observation emphasizes the need to elucidate molecular mechanisms that determine how MC linker flexibility and/or properties impact FtsH function and overall cellular physiology (53). Our work demonstrating the criticality of the FtsH MC linker in determining its function and has implications that potentially extend to all membrane-tethered proteins. We have shown that the molecular properties of the junction separating transmembrane and cytoplasmic domains of a protein can have profound impacts on its activity and anticipate that molecular hinges found in other membrane-proteins are key regulatory elements.

## Materials and Methods

### Strains and plasmids

*P. aeruginosa* strains used in this study were the clone C strain SG17M (41) and its derivative SG17M053 (SG17M Δ*ftsH1*Δ*ftsH2*) (5). For liquid growth, strains were cultivated in Luria-Bertani (LB) broth medium (BD Difco) at 37°C. If needed, 30 μg/mL gentamicin (for pJN105 plasmid selection in *P. aeruginosa* and *E. coli*) and 50 μg/mL kanamycin (for pET28a(+) expression plasmid selection in *E. coli*) were used. Plasmids used in this study are described in Table S3. *P. aeruginosa* SG17M genomic DNA was used as a template to amplify native Pa*ftsH1* and Pa*ftsH2* sequences for standard cloning into the expression vector pET-28a(+) (Novagen) between the NheI/XbaI restriction sites, upstream (5′) of the sequence encoding the in-frame 6xHis tag. For cloning and expression of wild-type and variant Pa*ftsH1* and Pa*ftsH2* proteins in *P. aeruginosa* SG17M, the L-arabinose inducible broad host-range expression vector pJN105 was used (54) (5). For the construction of Pa*ftsH* DNTR hybrid constructs (Constructs a-e; PaFtsH1^NFtsH2^; PaFtsH2^NFtsH1^), the desired regions of Pa*ftsH1* and Pa*ftsH2* were selectively amplified from *P. aeruginosa* SG17M genomic DNA, isolated, subjected to overlapping PCR to create custom hybrids and cloned into pJN105. Standard Gibson Assembly (NEB) cloning was used to construct linker variants PaFtsH2^H1-link-32^, PaFtsH2^H1-link-12^ and site-directed mutagenesis (QuickChange, Agilent) was used to create PaFtsH2^H1-link-10^, and all triplicate insertion MC linker PaFtsH variants. Some constructs were subcloned into pJN105, while others were directly cloned into pJN105.

All primer sequences used in this study are described in Table S4. Stable introduction of expression plasmids into *P. aeruginosa* SG17M Δ*ftsH1*Δ*ftsH2* was achieved by electroporation using Gene Pulser (BioRad) with 0.1 cm gap cuvettes operated at 13 kV/cm, 400 ohm and 25 μF. All constructs were confirmed by sequencing.

### Protein expression and purification

For protein induction, PaFtsH1-6xHis, PaFtsH2-6xHis, and PaFtsH2-6xHis variants were expressed from a pET28a(+) plasmid in *E. coli* strain BL21(DE3) (NEB) grown at 37°C in LB supplemented with 50 μg/mL kanamycin. At OD_600_ of 0.6, temperature was reduced to 30°C, and β-D-1-thiogalactopyranosidee (IPTG) was added to a final concentration of 0.5 mM in 1 L cultures and cells were harvested after 3 h. FtsH purification was performed as previously described with modifications (22, 55). Cell pellets were resuspended in Tris Buffered Saline (TBS) containing protease inhibitor (Pierce Protease Inhibitor Capsules, EDTA-free, ThermoFisher) and frozen. For protein purification, cell suspensions were pelleted by centrifugation and pellets were lysed by repeated freeze thaw cycles (4x) in which frozen cell pellets were thawed in a warm water bath and frozen in a dry ice and ethanol bath. The lysate was mixed at room temperature with 20 mL B-PER (ThermoFisher) containing 10 mg lysozyme, 5 µL Benzonase (Merck) and EDTA-free protease inhibitor for 20 min. Lysate was then centrifuged for 30 min at 30,000 RCF at 4°C, and supernatant was discarded. The resulting pellet containing cellular inclusion bodies was resuspended in resolubilization buffer (50 mM CAPS (pH 11.0), 2 mM MgSO_4_, 0.1 mM ZnCl_2_). After manual resuspension, ATP was added to a final concentration of 1 mM, followed by a dropwise addition of 15% *N*-lauroylsarcosine (Sigma) to a final concentration of 0.6%. The solution was mixed at room temperature for 30 min to allow for the resolubilization of inclusion bodies and then centrifuged for 30 min at 30,000 RCF and the supernatant was applied to a Ni Sepharose column (5 mL HisTrap HP; Cytiva) equilibrated in base buffer (50 mM Tris-HCl (pH 8.0), 100 mM NaCl, 20 mM imidazole, 5 mM MgSO_4_, 2 mM β-mercaptoethanol, 100 µM ZnCl_2_, 1 mM ATP) containing 0.6% *N*-lauroylsarcosine. The column was washed with 3 column volumes (CV) of the same buffer, and then subjected to a wash with 9 CV of buffer made of base buffer plus 0.5% NP-40 (note that *N*-lauroylsarcosine is omitted from this wash buffer to allow for protein refolding). The column was washed with 3 CV by a third and final solution composed of base buffer plus 10% glycerol, 0.1% NP-40, and 80 mM imidazole (the first step of a stepwise elution; we find FtsH is still bound to the column at this step). Finally, protein was eluted with 280 mM imidazole in base buffer containing 10% glycerol, and 0.1% NP-40. Fractions containing protein at >95% purity as judged by SDS-PAGE were combined, concentrated, and dialyzed overnight at 4°C in dialysis buffer (50 mM Tris-HCl (pH 8.0), 80 mM NaCl, 60 mM imidazole, 5 mM MgSO_4_, 2 mM β-mercaptoethanol, 10 µM ZnCl_2_, 10% glycerol). After dialysis, protein was concentrated, aliquoted, flash frozen in liquid nitrogen, and stored at −80°C.

For purification of radioactive proteins (^35^S-EcFtsH, ^35^S-PaFtsH1 and ^35^S-PaFtsH2), the same experimental procedure as above was followed with the following modifications to the protein induction conditions: *E. coli* BL21(DE3) (NEB) cells harboring a pET28a(+) expression plasmid encoding FtsH-6xHis were grown in 500 mL of M9 minimal media lacking methionine and cysteine at 37°C and induced at OD_600_ 0.6 with 0.5 mM IPTG. At the time of induction, 20 µCi/mL EasyTag Express ^35^S Protein Labeling Mix (PerkinElmer) was added to cultures and temperature was reduced to 30°C. After three hours of growth post-induction, cells were harvested via centrifugation and purification was carried out as described above.

Purification of Arc-st11-ssrA and λcI^N^ substrates were carried out as described (56, 57). For approximation of native molecular weights of purified FtsH enzymes, calibration curves were calculated by running molecular weight standards (BioRad cat. No. 151-1901) on a Superdex increase 3.2/3.0 analytical gel filtration column equilibrated in Buffer PD. Purified FtsH proteins were applied to the column and native molecular weights were calculated by fitting to a one-phase decay model.

### Growth assays

For assessing growth on solid media, wild-type *P. aeruginosa* SG17M and SG17M Δ*ftsH1*Δ*ftsH2* mutant harboring pJN105 vector control and derivaties with cloned FtsH variants were grown from a single colony into 10 mL LB medium overnight supplemented with 30 μg/mL gentamicin. The OD_600_ of the suspension was adjusted to 1 in LB medium and 10-fold serial dilution was made in a 96 well plate. 10 μL of the suspension was spotted on LB-agar plates with 30 μg/mL gentamicin and incubated at 37°C overnight. Colony size as a semiquantitative assessment of growth was monitored visually and imaged at different time points. For quantifying growth in liquid culture, bacteria were grown overnight in LB and resuspended to a final OD_600_ of 1.0 in phosphate-buffered saline (PBS). 1 µL suspension was added to 200 μL M63 minimal medium with the appropriate antibiotics in 96-well plates. Growth conditions were provided in a SpectraMax i3x (Molecular Devices), where the plate was incubated at 37°C under shaking for 24 h with OD_600_ measured every 30 min.

### Antibiotic susceptibility assays

*P. aeruginosa* clone C SG17M strains grown on LB-agar plates overnight were resuspended in PBS to a final OD_600_ of 0.1. An aliquot of this suspension was streaked out evenly on a Mueller Hinton Agar plate (containing 30 μg/mL gentamicin) to which a tobramycin disc (10 μg, Liofilchem) was subsequently placed at the center of the agar plate. The tobramycin-disc-containing plate was incubated at 37°C and the diameter of the inhibition zone was measured after 20-24 h.

### Enzymatic assays

Gel-based protein degradation assays were performed at 40 °C in Buffer PD (50 mM Tris-HCl (pH 8.0), 80 mM NaCl, 5 µM MgSO_4_, 12.5 µM ZnCl_2_, 10% glycerol, 2 mM β-mercaptoethanol, and 0.1% NP-40) supplemented with 5 mM ATP and an ATP regeneration system consisting of 50 µg/mL creatine kinase (Millapore-Sigma) and 5 mM creatine phosphate (Millipore-Sigma). Protein substrates were added to degradation reactions at the following concentrations: 40 mM β-casein (Sigma), 15 µM Arc-st11-ssrA, 15 µM λcI^N^-ssrA, and 15 µM untagged λcI^N^, respectively. For Arc-st11-ssrA degradation reaction, FtsH was added at a final concentration of 3.53 µM hexamer equivalents, and for reactions containing all other protein substrates, 3.04 µM FtsH hexamer equivalents were added. Reactions were quenched by the addition of SDS-loading buffer and boiled before separation by SDS-PAGE. Proteins in gels were stained with SYPRO orange (Sigma-Aldrich), and gels were imaged with a Typhoon FLA9500 scanner (GE Healthcare).

ATP-hydrolysis assays were performed at 40 °C using an NADH-coupled assay (58) containing 0.43 µM FtsH hexamer, 5 mM ATP, and NADH-coupled regeneration system in Buffer PD. ATP hydrolysis was assessed by the decrease in absorbance at 340 nm using a SpectraMax M5 plate reader (Molecular Devices).

### Peptide arrays

Arrays of 12-mer peptides were synthesized by standard Fmoc (9-fluorenylmethoxycarbonyl) solid-phase techniques and C-terminally linked to a cellulose membrane using a ResPep SL peptide synthesizer (Intavis). Arrays were incubated with gentle agitation in TBST (3 x 5 min) followed by overnight blocking in 5% BSA in TBST at 4°C. Blocked arrays were washed in TBST (2 x 5 min) followed by Buffer PD containing 0.03% NP-40 (2 x 5 min) at room temperature, and then with with fresh Buffer PD containing 0.03% NP-40, 1.25 mM ATPγS, 0.05% BSA, and 1 µM radioactive FtsH hexamer. After incubation with agitation for two hours at room temperature, the array was briefly washed with Buffer PD containing 0.03% NP-40 and 1 mM ATP (2 x 30 s), excess liquid from the blot was dried, and the dried blot was exposed to a Storage Phosphor Screen (Amersham) overnight at room temperature and imaged using a Typhoon FLA9500 scanner (GE Healthcare).

### Bioinformatic analyses

To computationally search for FtsH proteins across phyla, FtsH1 (accession number: EWH24232.1) and FtsH2 (accession number: EWH27927.1) of *P. aeruginosa* SG17M were used as queries to identify homologs in the NCBI and UniProt databases by BlastP using standard parameters (59, 60). Selection criteria were >93% coverage over the entire length of the amino acid sequence to account for the sequence diversity at the N- and C-termini of the protein combined with a lower identity limit of 50%. In addition, representative FtsH proteins, with the consideration of one representative protein per genus and up to three proteins per phylum, from all major bacterial phyla, representative proteins from other branches of the phylogenetic tree including plant FtsH representatives from *Arabidopsis thaliana* and mammalian representatives from *Homo sapiens* and selected protein sequences from UniProt were included in the assessment of phylogenetic relatedness. Subsequently, proteins were aligned using ClustalX2 using standard parameters (61). The aligned sequences were subjected to phylogenetic analysis using preliminary neighbor-joining (NJ) and conclusively maximum likelihood (ML) in MEGA7.0 (62). The Poisson model was used as an amino acid replacement model. The robustness of the phylogenetic tree topologies was evaluated by bootstrap analysis with 100 replications.

To analyze the linker sequences, the protein alignment was inspected manually in ClustalX2 (63) and GeneDoc (nrbsc.org/gfx/genedoc). Subsequently, proteins with linker sequences most homologous to the FtsH1 and FtsH2 linker sequence were selected with substantial similarity to the N-terminal alpha-helix sequence of FtsH1 and FtsH2 as a selection criterium. WebLogos which display the frequency of amino acid occurrence at one position were created using the resulting linker sequence alignments for FtsH1-like and FtsH2-like linkers, respectively (64).

Structural prediction of the MC linker in Figure 5C was calculated using PEP-FOLD 3.5 (65). Structural prediction of *P. aeruginosa* RpoH (PaRpoH) was calculated using ROSETTA (66). PaFtsH1 and PaFtsH2 monomer structures were predicted by AlphaFold (67) using accession numbers Q9HV38_PSEAE for PaFtsH1 and A0485GIM3_PSEI for PaFtsH2. The following residue changes were made for A0485GIM3_PSEI to achieve 100% sequence identity with PaFtsH2: G68S, I447M, and A524E in PyMOL 2.5.4 (The PyMOL Molecular Graphics System, Version 2.0 Schrödinger, LLC). Estimations of structural disorder were calculated by the IUPred algorithm (68). Charge versus hydropathy calculations were computed using PONDR® protein disorder predict (69–71).

### Electron microscopy and image processing

Samples were diluted to 0.4 mg/mL with storage buffer (50 mM Tris-HCl (pH 8.0), 10 mM KCl, 5 mM MgSO4, 10 µM ZnCl_2_, 60 mM imidazole, 10% glycerol, 0.1% Igepal and 2 mM β-mercaptoethanol). 3.0 µl of the dilutions were placed to glow-discharged copper grids with continuous carbon support film and incubated for 1 min. Excess sample was blotted away with filter paper and the grids washed in three drops of milli-Q water before staining 30 s in 2% uranyl acetate (TAAB Laboratories Equipment Ltd., UK). After most stain was blotted away with the filter paper, the grids were let to air-dry. Samples were checked in JEM-2100f field emission electron microscope (JEOL Ltd., Japan) operated at 200 kV. Images were collected at 50kx nominal magnification and 0.7-1.5 µm defocus with Tvips TemCam XF416-camera (Tietz Video and Image Processing Systems GmbH, Germany) using real-time drift correction.

All image processing was done in EMAN2 (version 2.2; (72). Particles were boxed in e2boxer_old using swarm mode using box size of 128 pixels. The extracted particles were corrected for the contrast transfer function. 2-D classifications were used for estimating data quality and resulting 2-D classes to create initial models both without symmetry and with the proposed C6-symmetry. Initial models were further refined during following 3-D refinement rounds. Final particle sets for 3D-reconstructions in e2refine_easy using C6 symmetry and defined target resolution 18.0Å contained 4781 (PaFtsH1_6_), 2921 (PaFtsH2_6_) and 3510 (PaFtsH2_6_^H1-link-32^) particles. Docking of atomic structures was achieved using UCSF Chimera (73).

## Data availability

All material, strains, and plasmids are available from either Tania A. Baker or Ute Römling.

## Acknowledgements

We thank Sanjay Hari and Sora Kim for generously providing materials and thoughtful discussions, and Igor Levchenko for the synthesis of peptide arrays. We thank Antonius Koller and Richard Schiavoni from Biopolymers & Proteomics Facility at the Koch Institute at MIT for conducting mass spectrometry. SMK received a personal scholarship from the Future University, Cairo, Egypt partially to conduct this work and travel grants from the Swedish Medical Association (SSMF) for a research stay at the Department of Biology, MIT, Cambridge, USA. This work was funded by the Swedish Research Council for Medicine and Health (2012-56X-22034-01-3).

## Author contributions

**GDM:** Conceptualization; Investigation; Methodology; Formal analysis; Writing – original draft; review & editing. **SMK:** Conceptualization; Investigation; Methodology; Formal analysis. **LC:** Investigation; Methodology; Formal analysis. **PP:** Investigation; Methodology; Formal analysis. **HH:** Supervision; Resources. **RTS:** Methodology; Resources; Writing – review & editing**. TAB:** Conceptualization; Formal analysis; Funding acquisition; Investigation; Methodology; Project administration; Resources; Supervision; Writing – original draft; Writing – review & editing. **UR:** Conceptualization; Data curation; Formal analysis; Funding acquisition; Investigation; Methodology; Project administration; Resources; Supervision; Writing – original draft; Writing – review & editing.

In addition to the <u>CRediT</u> author contributions listed above, the contributions in detail are:

Conceptualization: GDM, SMK, TAB, UR. Formal Analysis: GDM, SMK, LC, PP, HH, TAB, UR. Methodology: GDM, SMK, PP, LC, RTS, TAB, UR. Investigation: GDM, SMK, LC, PP, HH, UR. Resources: HH, RTS, TAB, UR. Manuscript writing: GDM, SMK, PP, RTS, TAB, UR with input from all authors.

## Supporting information

A PDF file with supporting information containing Figs. S1-S8 is provided.

