## Supplementary figures and images for "The membrane-cytoplasmic linker defines activity of FtsH proteases in *Pseudomonas aeruginosa* clone C"

### Supplemental Figures 1-8

#### Slide 1
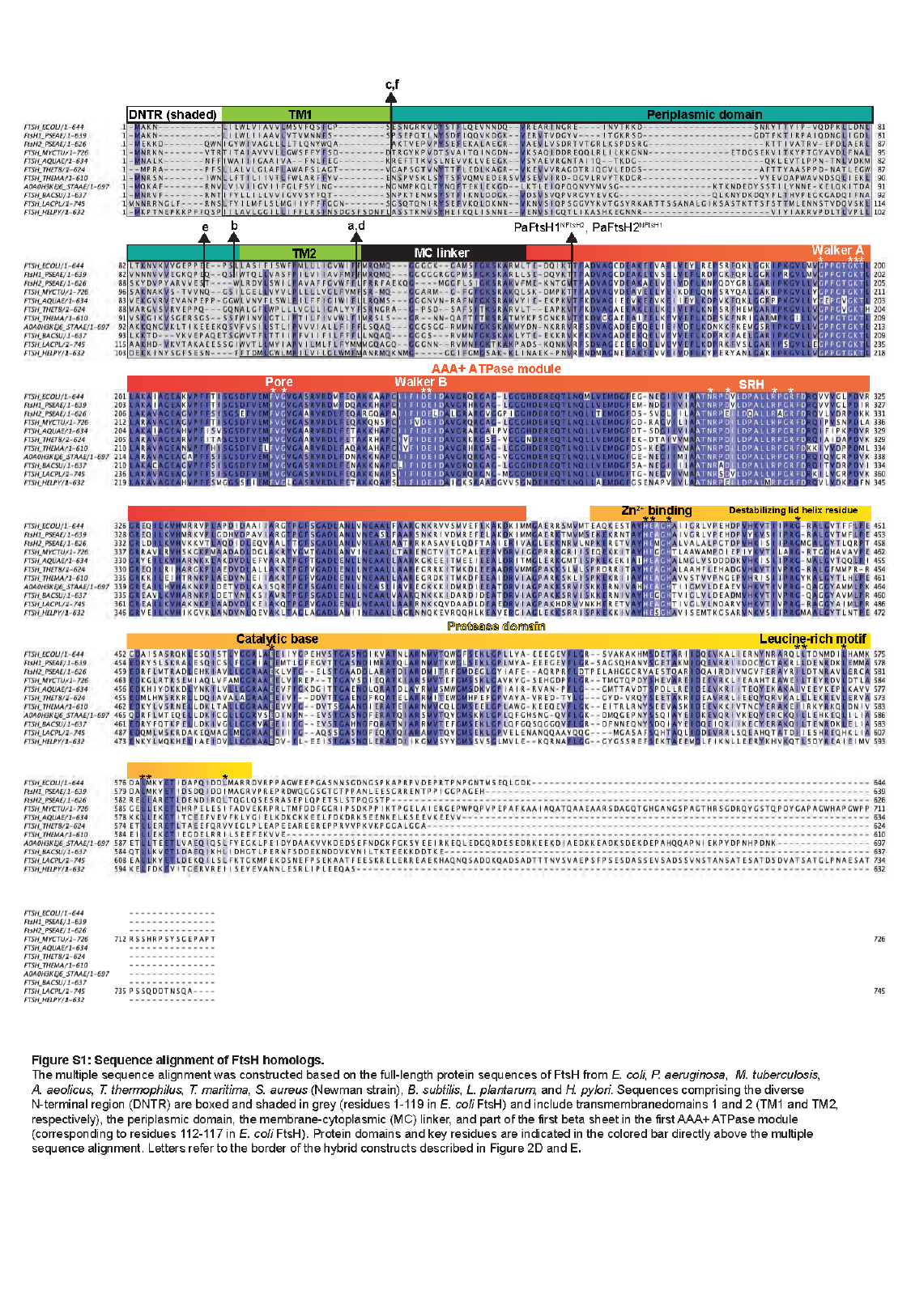

#### Slide 2
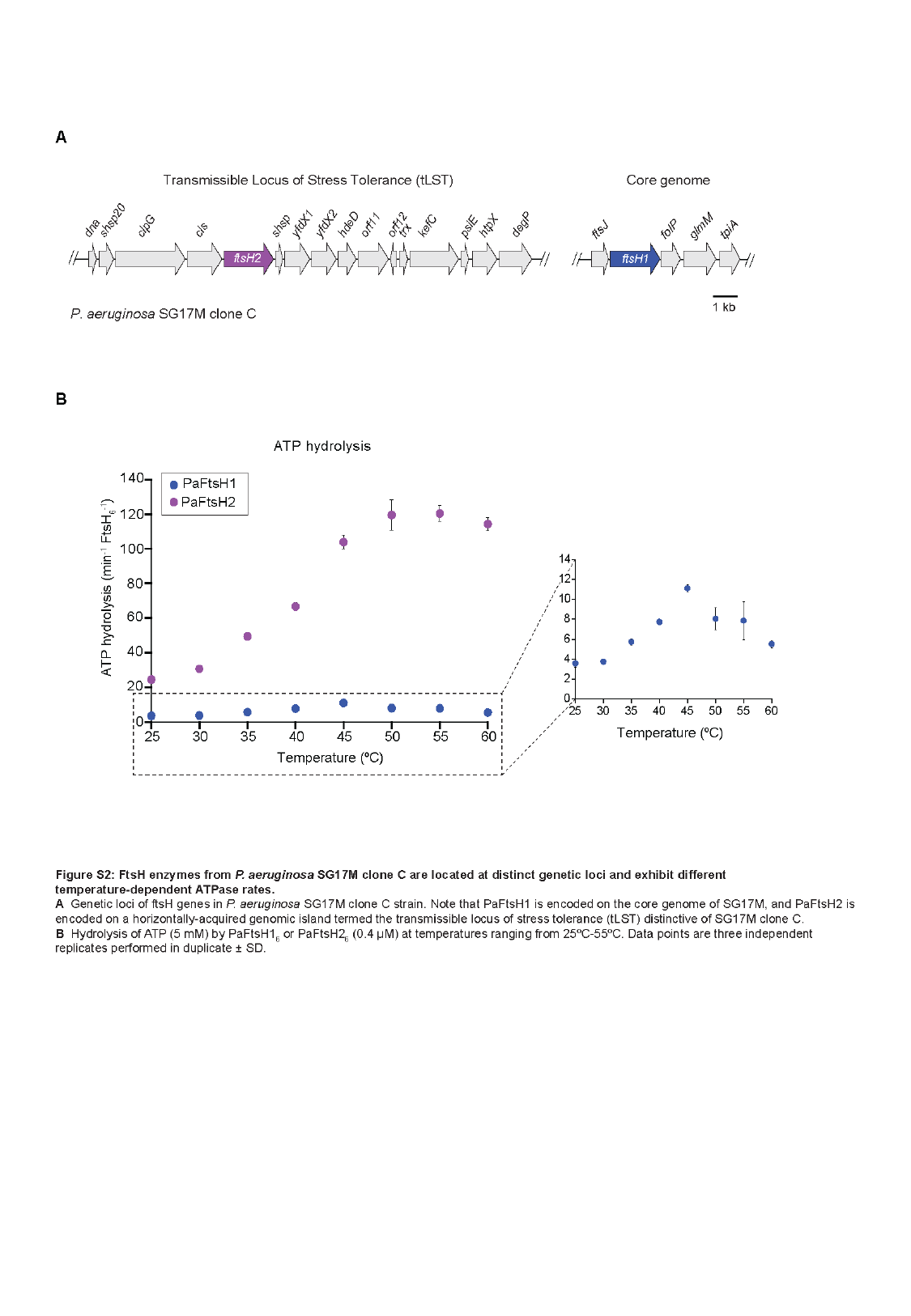

#### Slide 3
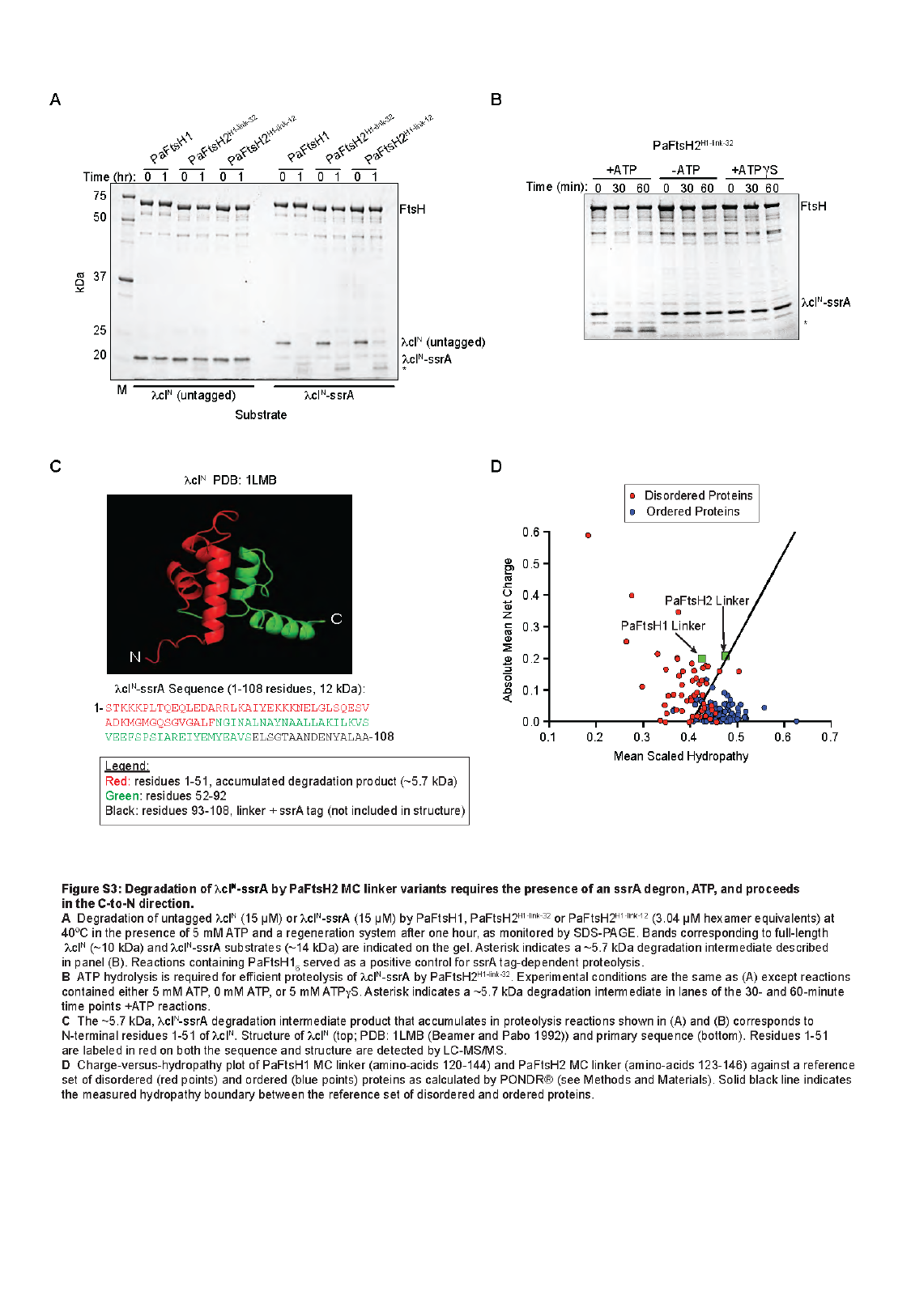

#### Slide 4
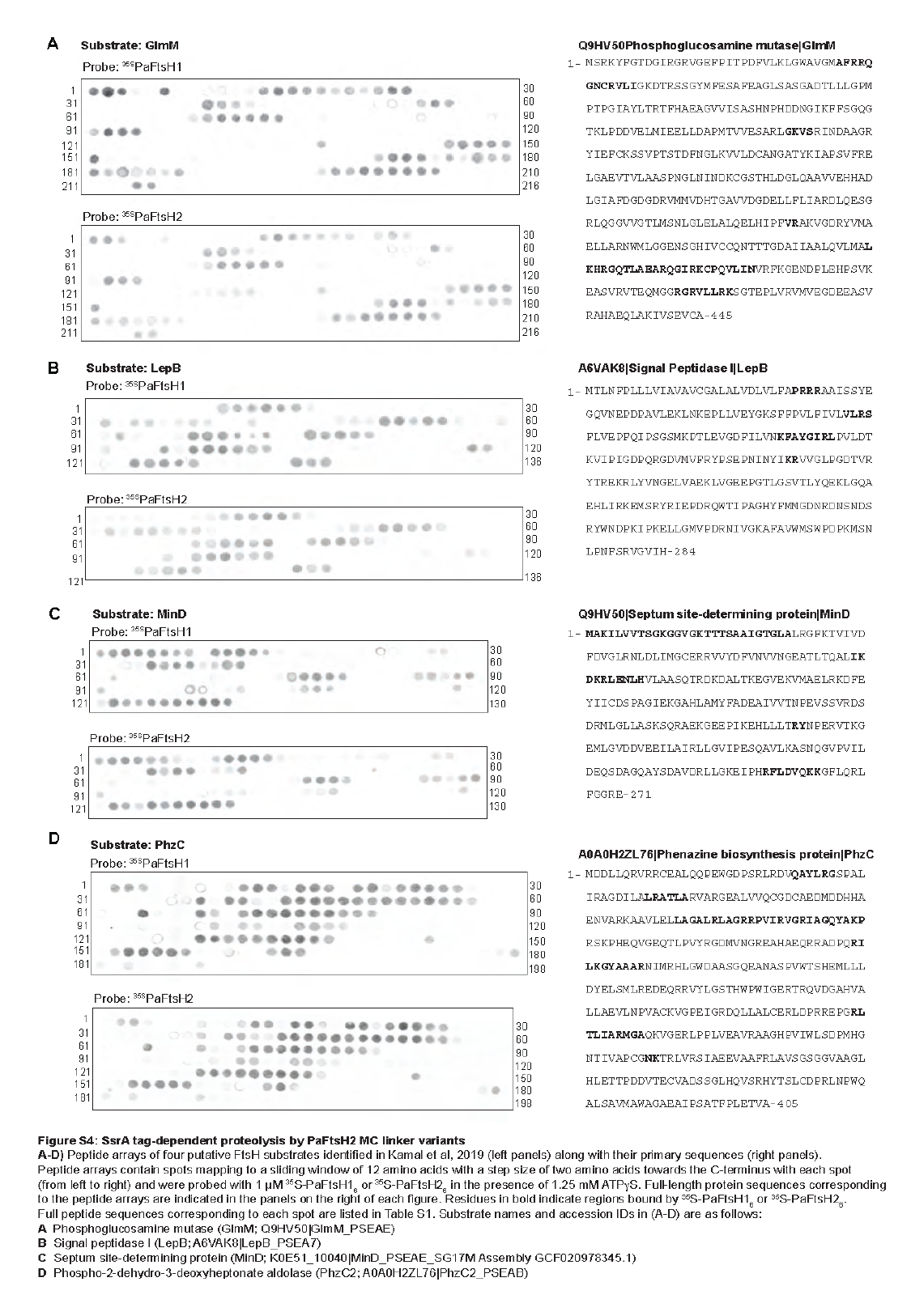

#### Slide 5
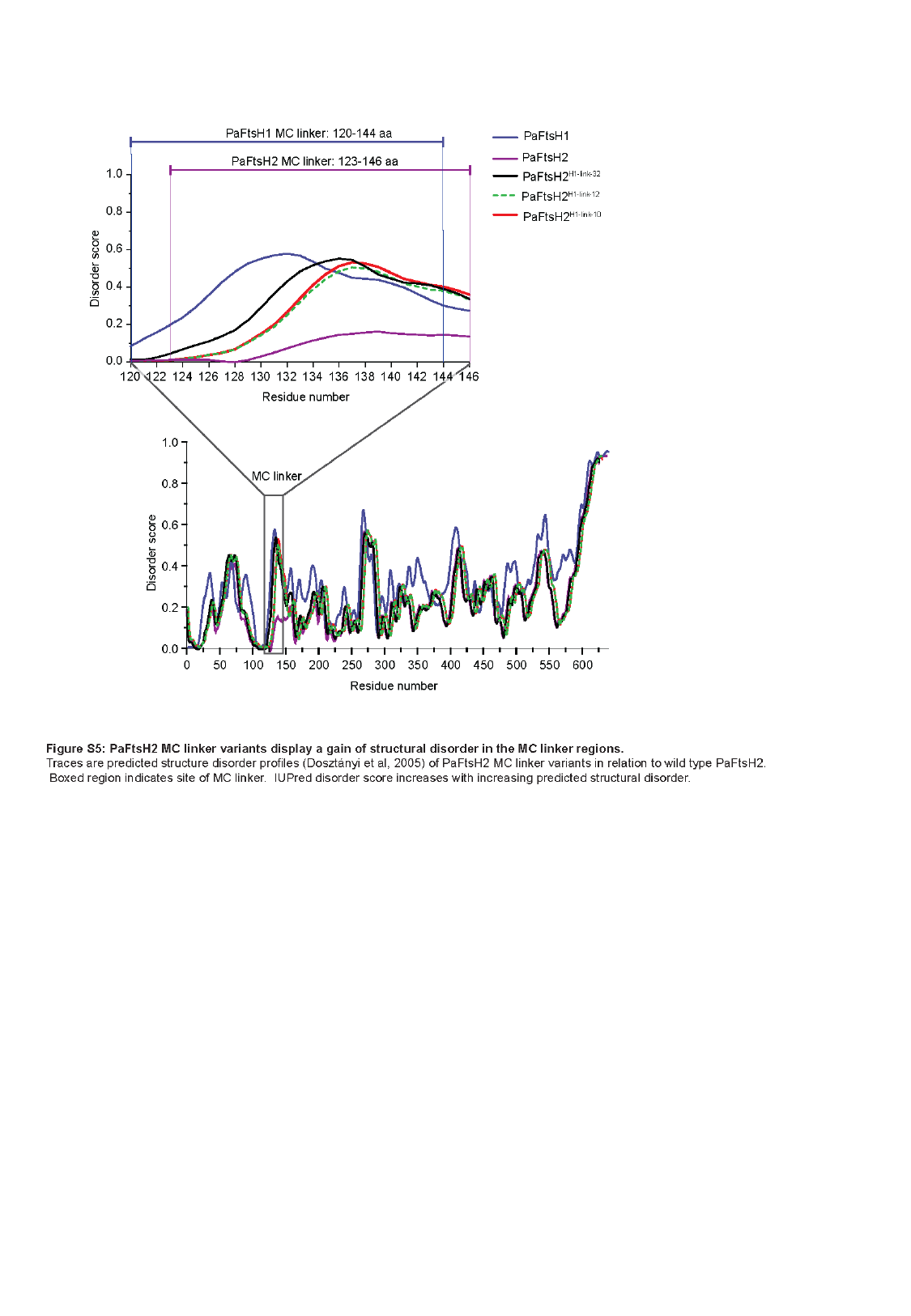

#### Slide 6
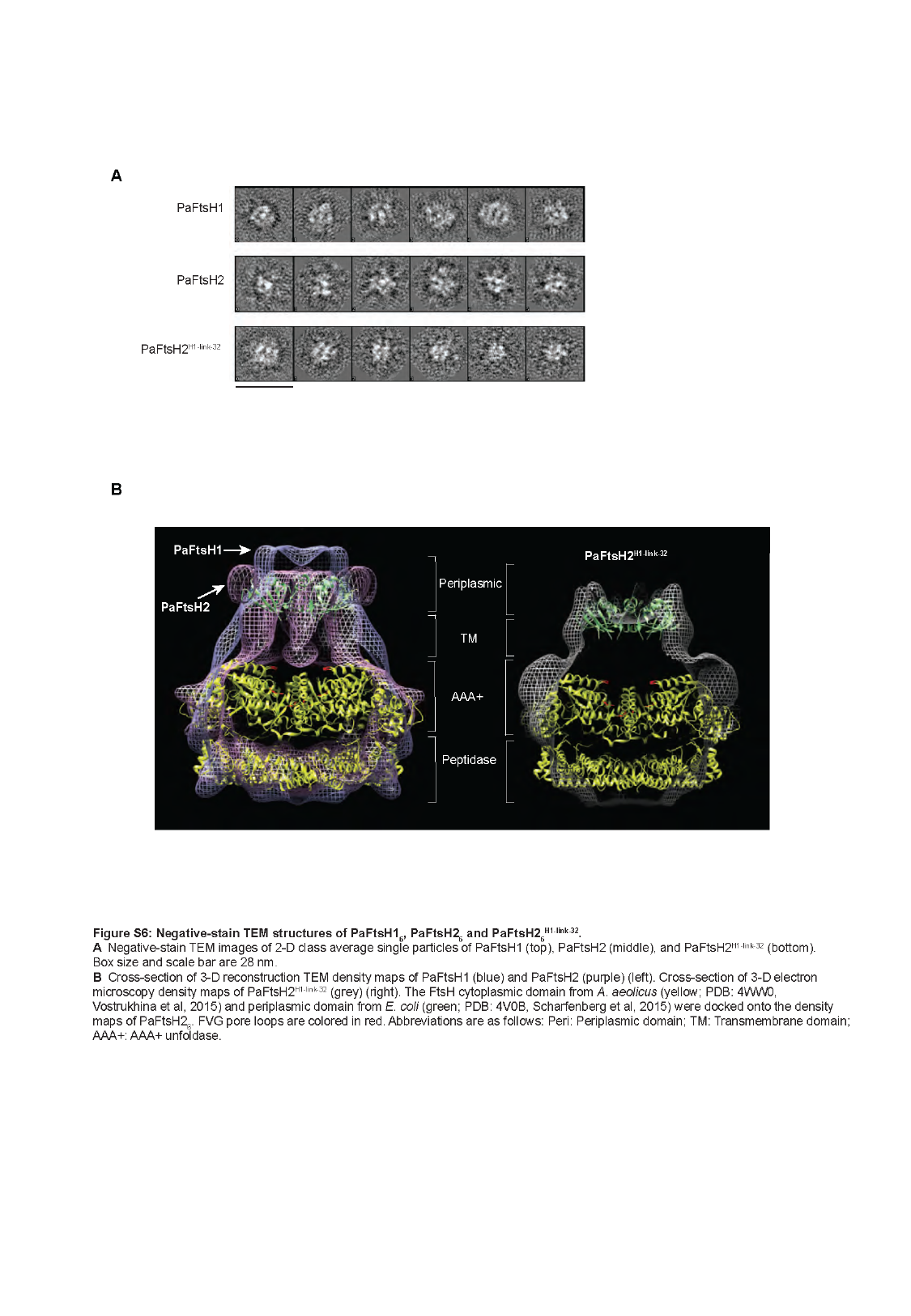

#### Slide 7
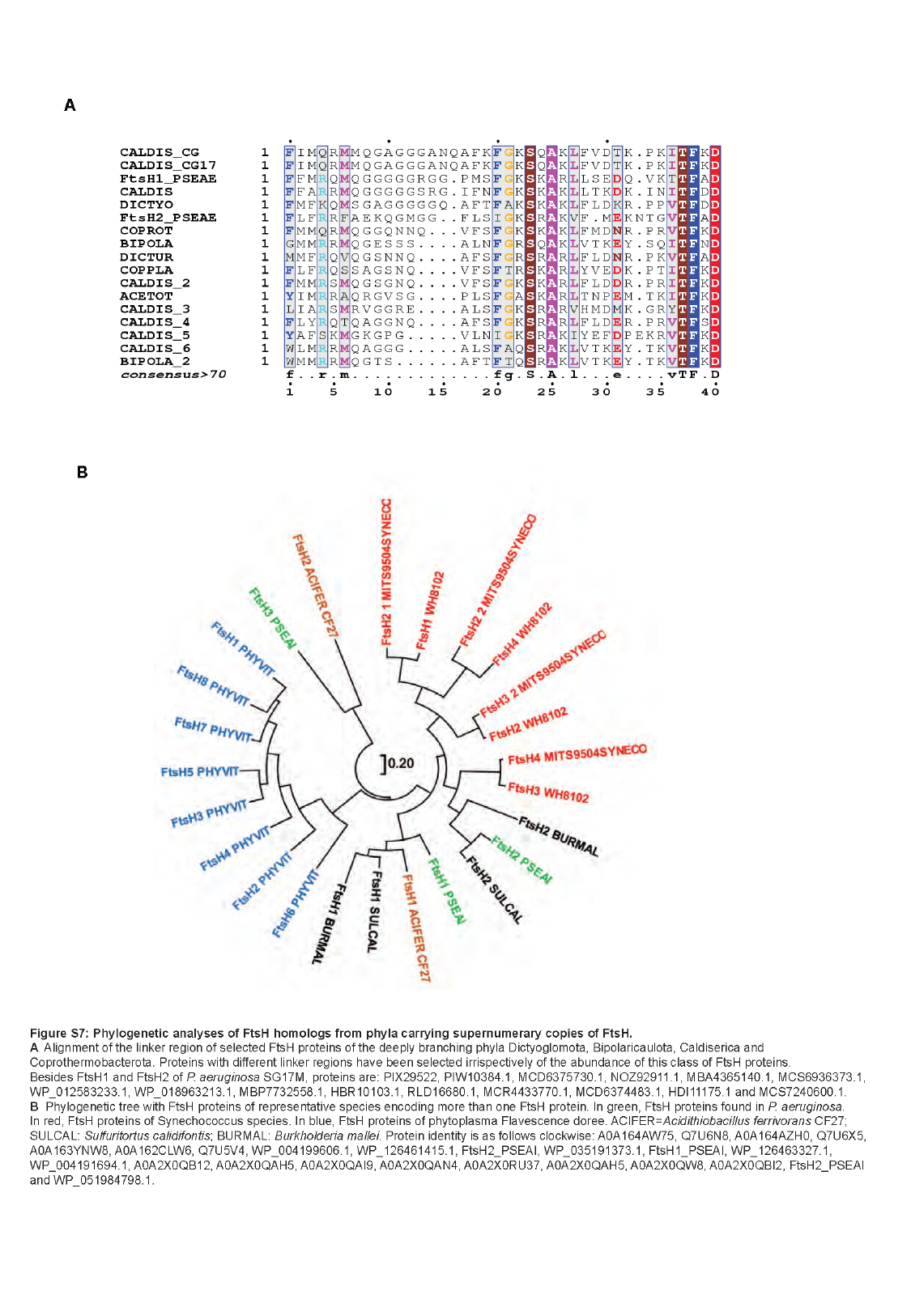

#### Slide 8
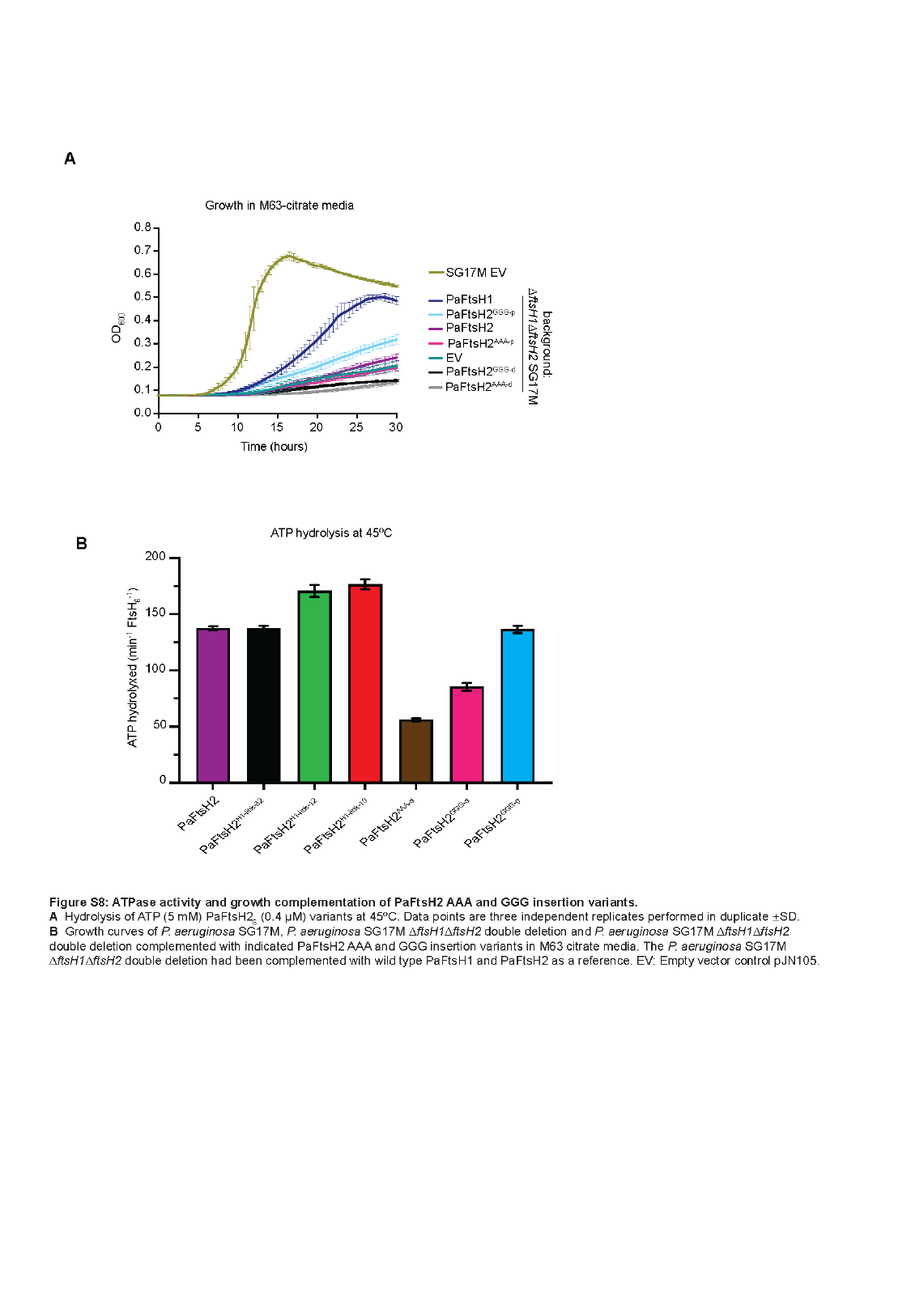
